# Decoupled recovery of soil physicochemistry, microbiome and methane sink function during secondary forest succession in the Eastern Himalayas

**DOI:** 10.64898/2026.10.04.753464

**Authors:** Mianhao Zhang, Xiaohui Zhao, Yuting Zhao, Dandan Shi, Wenzheng Fu, Cuiyun Yang, Hang Yu

## Abstract

Secondary forests are increasingly important for biodiversity recovery and ecosystem functioning after disturbance. As vegetation regenerates, accompanying shifts in soil conditions and microbial communities regulate climate-relevant processes such as atmospheric methane (CH_4_) uptake. Whether these belowground dimensions recover synchronously, however, remains unclear. Here we show contrasting patterns of belowground recovery across a forest chronosequence in Metok, Eastern Himalayas. Soil inorganic properties and microbial community composition converged progressively on the primary-forest state as soils acidified. By contrast, soil organic carbon, microbial diversity and abundance showed non-monotonic patterns across stand ages. Co-occurrence network topology resembled the primary forest by 40 years, yet the keystone taxa underlying it were entirely stage specific. Atmospheric CH_4_ uptake fluctuated: uptake in 25-year stands did not differ significantly from that in the primary-forest reference while uptake in 40-year stands was lower and comparable to that in the recently disturbed 5-year forest. The combined abundance of a taxonomically defined set of putative methanotrophs provided limited additional information for predicting CH_4_ up-take. Exploratory associations with individual amplicon sequence variants did not establish methane-oxidizing function or indicator value. Similarity to the primary-forest reference was not consistently greater in older secondary stands, cautioning against inferring restoration success from early succession alone and highlighting the need for longer chronosequences and repeated observation to resolve the dynamics of secondary-forest recovery.

## 1. Introduction

Forest succession refers to the temporal reorganization of ecosystem structure and eco-logical functioning following disturbance or environmental change (Horn, 1974; Chazdon, 2014). In human-modified landscapes, secondary succession has gained increasing ecological importance as land-use change has expanded the extent of regenerating secondary forests (Guariguata and Ostertag, 2001; Chazdon, 2014; Curtis et al., 2018). Understanding forest succession is therefore crucial for predicting the recovery of biodiversity, carbon sequestration and ecosystem services after disturbance (Chazdon et al., 2016; Poorter et al., 2021). Although forest recovery is commonly evaluated through aboveground attributes such as vegetation composition and biomass accumulation, belowground recovery represents an equally important dimension because soil biota and edaphic processes regulate biogeochemical cycling, carbon persistence, and greenhouse-gas exchange (Bardgett and Van Der Putten, 2014; Fierer, 2017). Therefore, examining belowground processes is essential for determining whether secondary succession restores ecosystem functioning rather than merely rebuilding aboveground forest structure.

Secondary succession after disturbance drives the concurrent reorganization of multiple belowground dimensions (Banning et al., 2011; Fierer, 2017; Jiang et al., 2021). However, recent studies increasingly suggest that these dimensions do not necessarily recover at the same rate or along the same trajectory (Liu et al., 2020; Poorter et al., 2021; Shen et al., 2025). Studies often examine microbial diversity, community composition, or individual ecosystem functions separately (Liu et al., 2021; He et al., 2022; Hu et al., 2022b; Beckstoffer et al., 2024), leaving uncertainty about whether microbial reassembly and ecosystem functioning recover in parallel (Sun and Badgley, 2019; Lin et al., 2026). Because soil physicochemical conditions also shift during succession and can independently regulate both microbial assembly and biogeochemical fluxes, abiotic, biotic, and functional recovery may proceed on different timelines (Zhang et al., 2022; Shen et al., 2025). Whether these belowground dimensions recover synchronously therefore remains an open question.

Soil greenhouse-gas fluxes provide a means of assessing whether abiotic, biotic, and functional recovery proceed synchronously during succession. Forest soils are major biological sinks for atmospheric methane (CH_4_), with a global synthesis estimating an annual uptake of approximately 15 Tg CH_4_ yr^−1^ (Feng et al., 2023). Soil CH_4_ uptake capacity can differ among forest successional stages (Liu et al., 2023), raising the question of whether CH_4_ sink strength recovers as forests regenerate. Forest disturbance can also increase soil carbon dioxide (CO_2_) efflux, with elevated emissions documented in logged tropical forests (Riutta et al., 2021). Although secondary forest regeneration promotes aboveground carbon accumulation (Chazdon et al., 2016), this does not necessarily imply a corresponding decline in soil CO_2_ efflux. It is therefore important to establish whether soil CO_2_ efflux decreases as forests regenerate. Both CH_4_ and CO_2_ fluxes are influenced by soil physicochemical conditions and biological processes. CH_4_ flux depends on bacterial and archaeal processes involved in methane oxidation and production (Meyer et al., 2020; Obregon Alvarez et al., 2023; Wigley et al., 2024), whereas soil CO_2_ efflux reflects contributions from both microbial and root respiration (Högberg et al., 2001). Because these controls can shift across succession, concurrent measurements of soil physicochemical conditions, microbial communities, and both gas fluxes along the same chronosequence are needed to assess whether CH_4_ sink recovery and changes in CO_2_ efflux parallel abiotic and biotic recovery.

Metok (Medog) County, located on the southern foothills of the Eastern Himalayas, offers a suitable setting for addressing this question. Situated within a biodiversity hotspot, the region encompasses pronounced elevational and hydroclimatic gradients, providing an opportunity to examine recovery in an environmentally heterogeneous mountain landscape (Myers et al., 2000; Hu et al., 2020; Wu et al., 2022). At the same time, historical slash-and-burn agriculture has created a mosaic of secondary forests of varying ages alongside remnant primary forest (Bryan et al., 2018; Wu et al., 2022), providing a basis for comparing abiotic, biotic, and functional recovery across successional stages. Recent studies in southeast Tibet have focused largely on high-elevation subalpine forests of the Sygera Mountains, typically between 3500 and 4300 m, and have shown that elevation, slope aspect, soil depth, and soil physicochemical properties are closely associated with microbial community structure (Li et al., 2022; Ma et al., 2022; Fu et al., 2023; Cheng et al., 2023; Ran et al., 2024). Although a recent study characterized elevational patterns in soil bacterial and fungal diversity and community composition along a 650–3800 m transect in Metok (Li et al., 2025), microbial recovery during secondary succession in low-elevation subtropical forests remains poorly understood. Beyond community recovery itself, a key question is whether soil CH_4_ sink function recovers in parallel. In reforested mine soils in the United States, microbial communities gradually converged to those in unmined reference forests, yet CH_4_ uptake capacity had not recovered after three decades (Sun et al., 2017; Sun and Badgley, 2019). These findings motivate a joint assessment of microbial community structure and soil CH_4_ uptake during secondary succession in Metok.

Here, we established a space-for-time chronosequence in Metok spanning 5-, 25-, and 40-year secondary forests, together with a primary-forest reference. We combined soil physico-chemical measurements, 16S rRNA gene amplicon sequencing, qPCR, and in situ CH_4_ and CO_2_ flux measurements with co-occurrence network analysis, differential-abundance analysis, and random forest regression. We addressed two questions: (i) do soil physicochemical properties, bacterial and archaeal communities, and CH_4_ uptake show concordant recovery patterns relative to the primary-forest reference; and (ii) which microbial taxa and environmental parameters best track CH_4_ sink function across forest secondary succession?

## 2. Materials and Methods

### 2.1. Study site, soil sampling and in situ CH_4_ and CO_2_ flux measurements

The study was conducted near Gelin Village, Metok County, Nyingchi, Tibet Autonomous Region, China (29^◦^13’N, 95^◦^11’E), on the southern foothills of the Himalayas. The region has a humid subtropical monsoon climate, with a mean annual temperature of 20–22 ^◦^C and annual precipitation of 2000–3000 mm. Native vegetation is montane subtropical evergreen broad-leaved forest. Historical slash-and-burn agriculture by the Menba ethnic group created heterogeneous disturbance across the landscape, and subsequent abandonment generated secondary forest stands at different stand ages.

To evaluate long-term natural recovery, we adopted a space-for-time chronosequence design (Fig. 1). Based on local land-use history, we selected four successional stages: 5-year herb–shrub establishment, 25-year pioneer tree dominance, 40-year secondary forest maturation, and a primary-forest (Fig. S1). Three independent plots per stage were established and served as plot-level replicates. All plots were located in comparable geomorphic units near Gelin Village, with similar slope and aspect, and were restricted to 1600–1850 m elevation to limit environmental confounding. Within each stage, replicate plots were separated by approximately 100 m in elevation to capture local topographic variability. Coordinates, elevation, and topographic attributes were recorded using a handheld GPS (Garmin, Olathe, KS, USA; Table S1).

**Figure 1:**
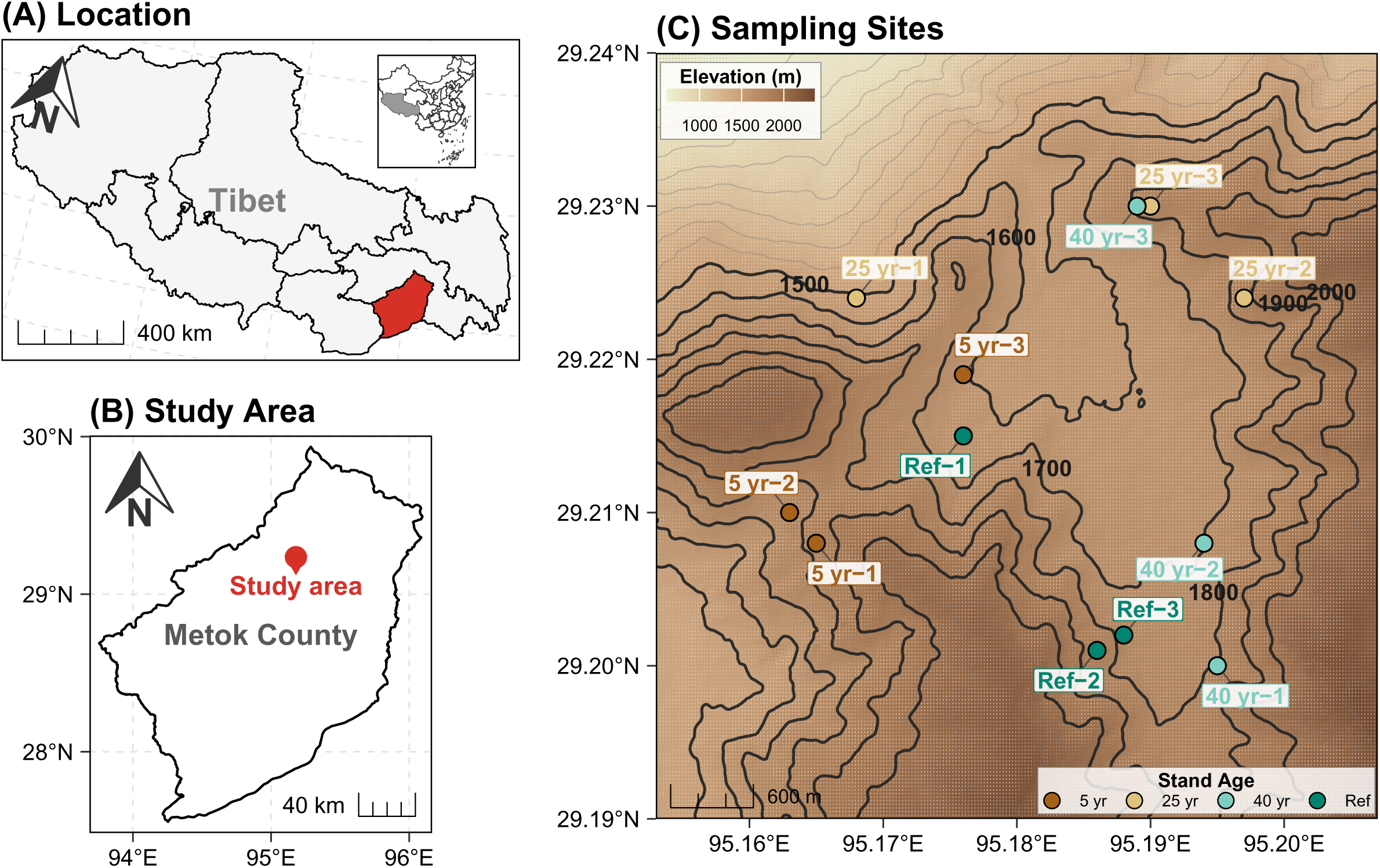
Geographic location of the study area and distribution of sampling plots in Metok County, Nyingchi, Tibet Autonomous Region, China. (A) Location of the study region within China. (B) Extent of the study area (approximately 28–30^◦^N, 94–96^◦^E). (C) Topographic map showing the distribution of sampling plots. The shaded background and contour lines indicate elevation, and colored circles represent sampling plots across successional stages.

Field measurements and soil sampling were conducted from 15 to 22 January 2025. Five sampling points were arranged within each plot using a five-point design, with at least 15 m between points. Sampling points were located at least 100 m from roads and anthropogenic edges to reduce edge effects.

Soil CH_4_ and CO_2_ fluxes were measured at each sampling point before soil coring. Measurements were conducted around local noon using an ultra-portable greenhouse-gas analyzer (LGR-915-0011, Los Gatos Research, San Jose, CA, USA) connected in a closed loop to a static chamber placed on a soil collar. Three consecutive measurements were taken, with the chamber automatically lifting and reclosing between measurements. Each cycle comprised 1 min of equilibration, 2 min of concentration recording, and 1 min of flushing/reset (Courtois et al., 2019). Only measurements with a linear concentration–time fit of *R*^2^ *>* 0.90 were retained for flux calculation. The three flux estimates for each gas were averaged to obtain a single value per sampling point. All CH_4_ fluxes were negative, indicating net atmospheric uptake, and their absolute values were used to express positive uptake rates. Positive CO_2_ fluxes indicated soil CO_2_ efflux.

Following flux measurements, surface litter was removed, and soil was collected from three locations arranged in an equilateral triangle within a 20 cm radius at each sampling point using a 5 cm-diameter stainless-steel auger. Surface (0–10 cm) and subsurface (10–20 cm) soils were sampled separately. After removing visible roots, stones, and coarse debris, the three cores from each depth were pooled into one composite sample, yielding 120 composite soil samples in total. Samples were placed in sterile polyethylene bags and transported in cool boxes to the field station on the day of collection, where they were stored at 4 ^◦^C. They were subsequently transported from the field station to the laboratory in cool boxes. In the laboratory, each soil sample was divided into two aliquots: one stored at *−*20 ^◦^C for DNA extraction and the other at 4 ^◦^C for physicochemical analyses.

### 2.2. Soil physicochemical analyses

For physicochemical analyses, fresh soil was reserved for water content (WC) determination, whereas the remaining aliquots were air-dried. Air-dried aliquots were passed through a 2-mm sieve for the determination of pH, electrical conductivity (EC), available phosphorus (AP), nitrate, and sulfate. Aliquots for total carbon (TC), total nitrogen (TN), and soil organic carbon (SOC) determination were finely ground and passed through a 250-*µ*m sieve. WC was determined gravimetrically by drying 5 g of fresh soil at 105 ^◦^C to constant weight and calculated as the mass lost during drying divided by the dry soil mass, expressed as a per-centage. Soil pH was measured potentiometrically in a 1:2.5 (w/v) soil-to-water suspension using a calibrated pH meter (Mettler Toledo, Switzerland). EC was measured in a 1:5 (w/v) soil-to-water extract using a conductivity meter (Mettler Toledo, Switzerland). AP was ex-tracted with 0.5 M NaHCO_3_ at pH 8.5 and determined by molybdenum–antimony colorimetry. After shaking and filtration, the filtrate was reacted with molybdate–antimony reagent, and absorbance was measured using a UV-2550 spectrophotometer (Shimadzu, Japan). AP concentrations were calculated against a KH_2_PO_4_ standard curve. For nitrate and sulfate determination, soil was mixed with water at a 1:5 (w/v) soil-to-water ratio and left to stand overnight. The suspensions were centrifuged at 5000 rpm for 10 min, and the supernatants were collected using a syringe and passed through 0.22 *µ*m membrane filters suitable for aqueous samples. Nitrate (reported as NO_3_^−^-N) and sulfate (SO_4_^2−^) in the resulting filtrates were quantified by ion chromatography (ICS-1100, Thermo Fisher Scientific, USA) using an anion-exchange column and an eluent containing 0.477 g L^−1^ NaHCO_3_ and 0.0672 g L^−1^ Na_2_CO_3_. TC and TN were measured in non-acidified soil by high-temperature dry combustion using an elemental analyzer (Vario EL III, Elementar Analysensysteme GmbH, Hanau, Germany). For SOC determination, approximately 50 mg of finely ground soil was weighed into silver capsules, moistened with 5% H_3_PO_4_ to remove inorganic carbon, and redried. SOC was then measured by dry combustion using an elemental analyzer (FlashSmart, Thermo Fisher Scientific, Waltham, MA, USA). The soil C:N ratio was calculated as SOC/TN. Elemental and ion concentrations were expressed on a 105 ^◦^C oven-dry soil mass basis.

### 2.3. DNA extraction, PCR amplification, and amplicon sequencing

Total genomic DNA was extracted from 0.25 g of soil using the DNeasy PowerSoil Kit (QI-AGEN, Hilden, Germany) according to the manufacturer’s instructions. DNA concentration was quantified with the Qubit dsDNA Assay Kit (Thermo Fisher Scientific, USA). For bacterial and archaeal community profiling, the V4–V5 region of the 16S rRNA gene was amplified using the archaeal/bacterial primer pair 515F (5’-GTGYCAGCMGCCGCGGTAA-3’) and 926R (5’-CCGYCAATTYMTTTRAGTTT-3’) (Parada et al., 2016), with Illumina adapters attached to the 5’ ends of the primers. PCR amplification was carried out in triplicate 25 *µ*L reactions containing 12.5 *µ*L of 2*×* KAPA HiFi HotStart ReadyMix (KAPA Biosystems), 0.2 *µ*M of each primer, and approximately 10 ng of template DNA. The amplification program consisted of an initial denaturation at 95 ^◦^C for 3 min, followed by 25 cycles of 98 ^◦^C for 20 s, 55 ^◦^C for 15 s, and 72 ^◦^C for 30 s, with a final extension at 72 ^◦^C for 5 min. Triplicate PCR products from the same composite soil sample were then combined, purified using AMPure XP beads (Beckman Coulter, USA), quantified again by Qubit, and pooled in equimolar amounts for library preparation. Paired-end sequencing (2 *×* 250 bp) was performed on an Illumina NovaSeq 6000 platform (Magigene Biotechnology Co., Ltd., Guangzhou, China).

### 2.4. Quantification of bacterial 16S rRNA, archaeal 16S rRNA, and fungal ITS gene abundance by qPCR

Total prokaryotic (bacterial and archaeal) 16S rRNA gene abundance was quantified by real-time quantitative PCR (qPCR) using the primer pair 515F (5’-GTGYCAGCMGCCGCGGTAA-3’) and 806R (5’-GGACTACNVGGGTWTCTAAT-3’) (Walters et al., 2015). Archaeal 16S rRNA gene abundance was quantified separately using the primer pair 344F (5’-ACGGGGCGCAGCAGGCGCGA-3’) and 517R (5’-ATTACCGCGGCTGCTGG-3’) (Bano et al., 2004). Fungal ITS abundance was quantified by targeting the internal transcribed spacer (ITS) region using the primer pair ITS1F (5’-CTTGGTCATTTAGAGGAAGTAA-3’) and ITS2 (5’-GCTGCGTTCTTCATCGATGC-3’) (White et al., 1990; Gardes and Bruns, 1993). All qPCR assays were run in triplicate on a QuantStudio 5 Real-Time PCR System. Reactions were performed in 20 *µ*L volumes containing 10 *µ*L of SYBR Green Master Mix and 0.4 *µ*M of each primer. Approximately 10 ng of template DNA was used per sample reaction. For the total prokaryotic and archaeal 16S rRNA gene assays, the thermal program consisted of an initial denaturation at 95 ^◦^C for 10 min, followed by 40 cycles of 95 ^◦^C for 30 s, 55 ^◦^C for 1 min, and 72 ^◦^C for 1 min. For the fungal ITS assay, amplification consisted of 40 cycles of 95 ^◦^C for 10 s, 60 ^◦^C for 30 s, and 72 ^◦^C for 30 s. Product specificity was assessed by melting-curve analysis from 65 to 95 ^◦^C with 0.5 ^◦^C increments. Reactions showing non-specific amplification were excluded.

Commercial pTOPO-T plasmids carrying cloned bacterial 16S rRNA gene, archaeal 16S rRNA gene, and fungal ITS fragments were obtained (Meiyou Anuo Biotechnology Co., Ltd., Beijing, China) and used as standards. For standard plasmid construction, the bacterial 16S rRNA gene fragment was amplified using primers 27F (5’-AGAGTTTGATYMTGGCTC-3’) and 1492R (5’-GGYTACCTTGTTACGACTT-3’) (Trembath-Reichert et al., 2016), and the archaeal 16S rRNA gene fragment was amplified using primers Arc23F (5’-TTCCGGTTGATCCYGCCGGA-3’) and U1492R (5’-GGYTACCTTGTTACGACTT-3’) (Scheller et al., 2016). The fungal ITS fragment was amplified using primers ITS1-modified (5’-TCCGTWGGTGAACCWGCGG-3’) (Babusci and Gunsch, 2026) and ITS2 (5’-GCTGCGTTCTTCATCGATGC-3’) (White et al., 1990). Before use as standards, plasmids were linearized with HindIII and purified (Bustin et al., 2009). Standard curves were generated separately for each assay using serial dilutions of the corresponding plasmids. Amplification efficiencies were 102.74%, 102.37%, and 101.95% for the total prokaryotic 16S rRNA gene, archaeal 16S rRNA gene, and fungal ITS assays, respectively, with *R*^2^ = 0.99 for all three standard curves. Gene copy numbers were calculated from the corresponding standard curves and expressed as copies g^−1^ dry soil. For each sample, bacterial 16S rRNA gene abundance was estimated by subtracting archaeal gene copy numbers from total prokaryotic gene copy numbers on the same dry-soil basis. Subtraction was performed on untransformed copy numbers before log_10_ transformation for statistical analyses.

### 2.5. 16S rRNA gene sequence processing and bioinformatics analysis

Paired-end reads from 16S rRNA gene V4–V5 amplicon sequencing were processed to generate an amplicon sequence variant (ASV) abundance table and assign taxonomy for bacterial and archaeal community analyses. Reads were demultiplexed using sample-specific barcodes and analyzed in QIIME 2 (v2024.2) (Bolyen et al., 2019). Primer sequences were removed using the Cutadapt plugin with the 515F and 926R primer sequences. Quality filtering, denoising, paired-read merging, and chimera removal were performed using the q2-dada2 denoise-paired method (Callahan et al., 2016). Forward and reverse reads were truncated to 230 and 210 bp, respectively, with no additional 5^′^ trimming. The maximum expected error was set to 2 for each read direction, the quality truncation threshold to 2, and the minimum overlap for paired-read merging to 12 bp. Samples were denoised independently, and chimeras were removed using the consensus method; other parameters were retained at their default values.

Representative ASV sequences were taxonomically assigned using the Naive Bayes classify-sklearn classifier (Bokulich et al., 2018), trained on the 515F–926R region extracted from the SILVA release 138 reference sequences clustered at 99% identity (Quast et al., 2013), with a confidence threshold of 0.8. ASVs unclassified at the kingdom level or assigned to mitochondria or chloroplasts were removed. The filtered dataset contained 8,944,341 sequences across 120 samples (mean: 74,536; range: 34,563–212,272), representing 68,920 ASVs in total.

### 2.6. Statistical analysis

All statistical analyses and visualizations were performed in R (v4.3.2; R Core Team, 2023). Soil physicochemical properties and CH_4_ and CO_2_ fluxes were analyzed using linear mixed-effects models. Fixed effects included successional stage, soil depth, elevation, and their interaction for soil properties, and successional stage and elevation for gas fluxes. For CH_4_ uptake, an additional model included WC as a covariate to assess whether differences among successional stages persisted after adjustment for WC. Both model types included random intercepts for sampling points nested within plots. Nonsignificant interaction terms were excluded from the models used for inference. Pairwise comparisons among stages were based on estimated marginal means, with *P* values adjusted for multiple comparisons. Model residuals were assessed for normality and homogeneity of variance, and variables were transformed when necessary.

Bacterial and archaeal community composition was analyzed using Bray–Curtis dissimilarities. Homogeneity of multivariate dispersion was assessed using betadisper. Associations of community composition with successional stage, soil depth, elevation, and soil physicochemical variables were evaluated using marginal PERMANOVA (adonis2; 999 permutations). Permutations were restricted to reflect the sampling hierarchy of points nested within plots and plots nested within stages, with plots as the highest independent units for stage-level inference. Constrained community variation was visualized using distance-based redundancy analysis (db-RDA), with significant environmental vectors fitted to show associations between community turnover and soil physicochemical gradients.

Compositional recovery was quantified separately for bacteria and archaea using Bray–Curtis distances to the primary-forest reference, calculated from relative-abundance ASV matrices. All comparisons were restricted to samples from the same soil depth. The dissimilarity between composite soil sample *i* and primary-forest sample *j* was calculated as

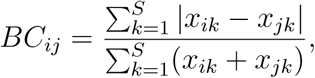

where *x_ik_* and *x_jk_* are the relative abundances of ASV *k* in samples *i* and *j*, respectively, and *S* is the total number of ASVs. For each secondary-forest sample *i* at depth *d*, distance to the reference was calculated as

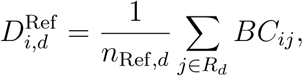

where *R_d_* is the set of all primary-forest samples at depth *d* and *n*_Ref*,d*_ is its size. Within-reference distances were calculated by excluding the focal primary-forest sample *r*:

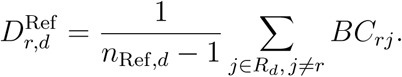

Lower values indicate greater compositional similarity to the primary-forest reference.

Associations between microbial recovery metrics and soil physicochemical variables were assessed using separate linear mixed-effects models for each response–predictor pair. Responses included bacterial and archaeal Bray–Curtis distances to the primary-forest reference, bacterial and archaeal Shannon diversity calculated from rarefied ASV tables, and log_10_-transformed bacterial and archaeal 16S rRNA and fungal ITS gene abundances. Predictors included pH, water content, total carbon, soil organic carbon, total nitrogen, soil C:N ratio, available phosphorus, nitrate, sulfate, and electrical conductivity. For each pair, the microbial response and soil predictor were standardized to *z* scores before model fitting. Models included soil depth, elevation, and successional stage as fixed covariates and random intercepts for sampling points nested within plots. Significance was assessed from the coefficient of the standardized soil predictor, with *P* values adjusted across tests using the Benjamini–Hochberg false-discovery-rate (FDR) procedure.

Bacterial and archaeal co-occurrence networks were constructed for each successional stage using ggClusterNet (v0.1.0) (Wen et al., 2022). Rare taxa were removed using pre-defined relative-abundance (*<* 0.01%) and prevalence (*<* 20% of composite soil samples) thresholds in all stages. Pairwise associations in the compositional data were estimated using SparCC (Friedman and Alm, 2012), and their significance was assessed using 1000 bootstrap resamples. Edges with *|r| >* 0.6 and FDR-adjusted *P <* 0.05 were retained in undirected networks; both positive and negative associations were included unless otherwise stated. Topological properties, including node and edge counts, mean degree, and modularity, were calculated using igraph (Csardi and Nepusz, 2006). Modules were detected using the fast-greedy algorithm (`cluster_fast_greedy`). Nodes were classified by within-module connectivity (*Z_i_*) and among-module connectivity (*P_i_*) as peripherals (*Z_i_ ≤* 2.5, *P_i_ ≤* 0.62), connectors (*Z_i_ ≤* 2.5, *P_i_ >* 0.62), module hubs (*Z_i_ >* 2.5, *P_i_ ≤* 0.62), or network hubs (*Z_i_ >* 2.5, *P_i_ >* 0.62) (Guimerà and Nunes Amaral, 2005).

Differentially abundant taxa across successional stages were identified from raw ASV counts using ANCOM-BC2 (ANCOMBC v2.4.0) (Lin and Peddada, 2024). Successional stage was the main fixed effect, with primary forest as the reference level; soil depth, elevation, and retained environmental covariates were included as additional fixed effects. Taxa were considered significant when the Holm-adjusted *q* value was *<* 0.05 and the sensitivity indicator (`diff_robust_abn`) was TRUE. Bias-corrected log-fold changes (LFCs) relative to primary forest were used for visualization. For heatmaps, LFCs were re-centered within each key ASV relative to the stage in which it was identified as a keystone. Associations between keystone ASV relative abundances and soil physicochemical variables were assessed using Spearman’s rank correlation, with coefficients reported as *ρ* and *P* values adjusted using the Benjamini–Hochberg procedure.

Putative methane-cycling ASVs were identified from SILVA-based taxonomic assignments. The putative aerobic methanotroph set comprised 61 ASVs assigned to nine genera with reported methanotrophic representatives: *Methylobacter*, *Methylomonas*, *Methylocystis* and *Methylosinus* (Bowman et al., 1993); *Methylocella* (Dedysh et al., 2000); *Methylosoma* (Rahalkar et al., 2007); *Methyloferula* (Vorobev et al., 2011); *Candidatus* Methylospira (Danilova et al., 2016); and *Methylocapsa* (Tveit et al., 2019). Membership was defined independently of ASV–CH_4_ flux associations. For this bacterial set, combined relative abundance was calculated as *g/b*, where *g* is the summed read count of the selected ASVs and *b* is the total bacterial read count. For modelling, this proportion was expressed as ln[(*g* + 0.5)*/*(*b − g* + 0.5)]. Group-associated 16S rRNA gene copy abundance was estimated as [(*g* + 0.5)*/*(*b* + 1)]*B* and log_10_-transformed, where *B* is the bacterial 16S rRNA gene copy abundance estimated from qPCR, expressed as copies g^−1^ dry soil. This was an indirect estimate of group-associated 16S abundance. Putative methanogenic archaea were identified separately as ASVs assigned to known methanogenic lineages, including Methanomassiliicoccales, Methanomethyliaceae, and *Methanosarcina*. Their relative abundances and within-group taxonomic contributions were calculated from the rarefied ASV table and summarized by successional stage.

Random forest regression models were fitted using ranger (v0.16.0) to identify microbial and edaphic predictors of CH_4_ and CO_2_ fluxes (Wright and Ziegler, 2017). For microbial models, ASVs present in *>* 20% of composite soil samples were retained, and their counts were centered log-ratio (CLR) transformed with a pseudocount of 1 (Aitchison, 1982). Edaphic models used measured soil physicochemical variables directly. All models were fitted with 2000 trees, and variable importance was assessed by permutation. Hyperparameters, including mtry, `min.node.size`, and `sample.fraction`, were optimized by repeated cross-validation grouped by plot to prevent information leakage between observations from the same plot. Model performance was evaluated using cross-validated *R*^2^, RMSE, and MAE. Predictors ranked among the top variables in *>* 60% of repeated iterations were considered stable; the top 20 ASVs were used to define the stable predictor set for the microbial CH_4_ model. CH_4_-associated key ASVs were defined as stable microbial predictors that were also significantly differentially abundant among stages according to ANCOM-BC2 and classified as network connectors or module hubs. The same intersection procedure was applied to CO_2_ flux.

Associations between log_10_-transformed relative abundances of CH_4_-associated keystone ASVs and CH_4_ uptake rates were assessed using Pearson’s correlation and simple linear regression. Uptake rates were defined as the magnitudes of measured negative CH_4_ fluxes. Pearson’s *R* and corresponding *P* values were reported. The same analyses were used to screen succession-associated keystone ASVs for associations with measured CO_2_ efflux.

Representative 16S rRNA gene V4–V5 sequences of CH_4_-associated *Mycobacterium* ASVs were compared with published methane-associated *Mycobacterium* references using BLASTn against the NCBI nucleotide database. For each ASV, top-hit identities, percent identity, alignment length, and query coverage were recorded.

## 3. Results

### 3.1. Edaphic variation across successional stages

The chronosequence near Gelin Village in Metok, Eastern Himalayas, encompassed forest regeneration following slash-and-burn agriculture, from herb–shrub establishment at 5 years to pioneer tree dominance at 25 years and secondary forest maturation at 40 years (Fig. 1; Fig. S1). To assess soil abiotic recovery alongside this vegetation recovery, we compared soil physicochemical properties across the three secondary-forest stages and a primary-forest reference in surface (0–10 cm) and subsurface (10–20 cm) soils. These comparisons revealed two main patterns: directional changes toward the primary-forest reference and stage-specific fluctuations (Fig. 2; Table S2). Most properties also varied significantly with soil depth, although their patterns across succession were broadly similar between surface and subsurface soils (Fig. S2; Table S2).

**Figure 2:**
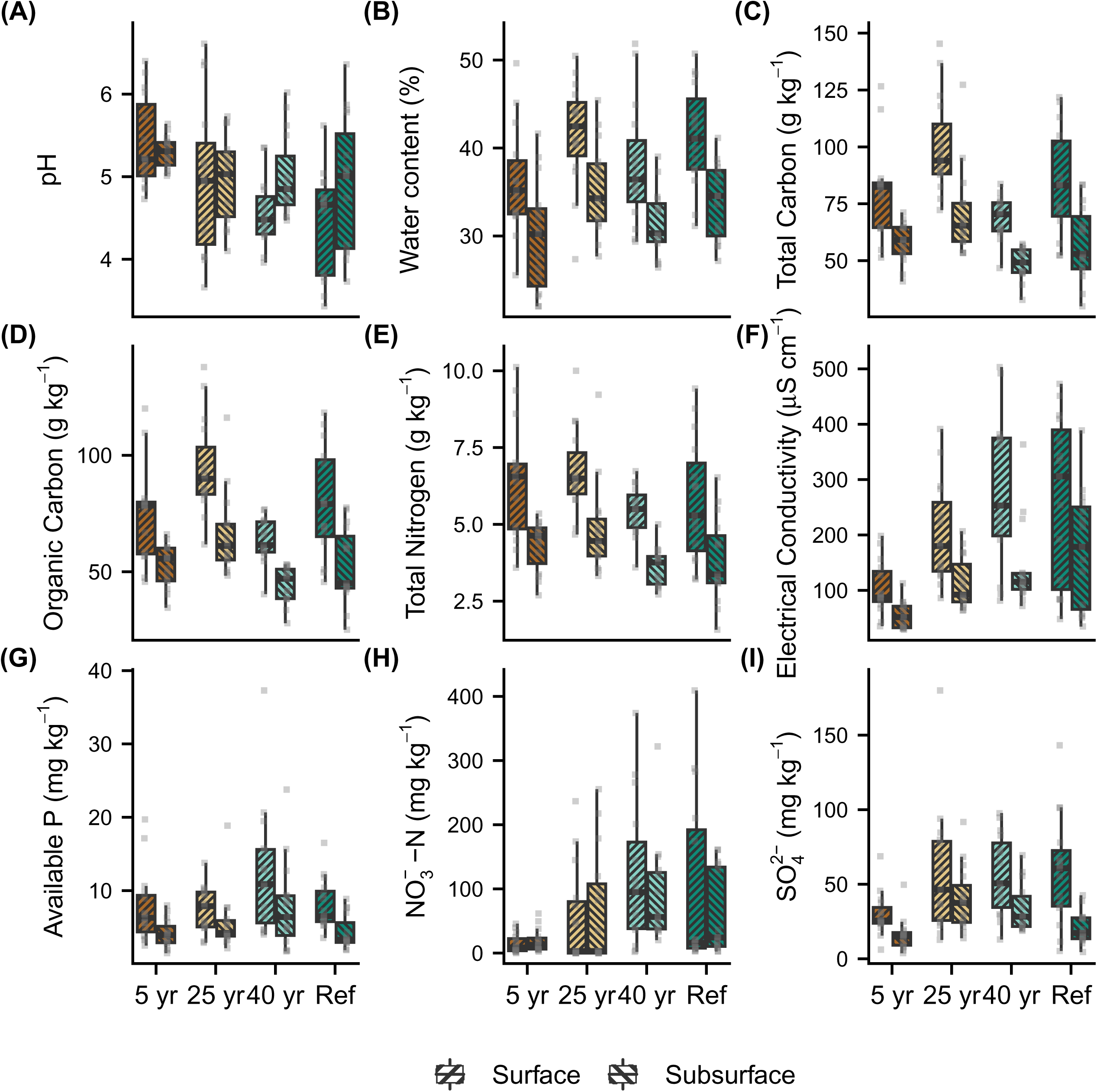
Soil physicochemical properties among successional stages and soil depth. Panels show (A) pH, (B) soil water content, (C) total carbon, (D) soil organic carbon, (E) total nitrogen, (F) electrical conductivity, (G) available phosphorus, (H) NO_3_^−^-N, and (I) SO_4_^2−^. In each boxplot, boxes represent 25th–75th percentiles of the data distribution with whiskers at 1.5 times the interquartile range and the middle line representing median. Points represent composite soil samples (*n* = 15 per stage and depth; five sampling points in each of three plots). Surface and Subsurface denote 0–10 and 10–20 cm, respectively.

Soil pH, electrical conductivity (EC), NO_3_^−^-N, and SO_4_^2−^ showed directional changes toward the primary-forest reference. Soil pH decreased from 5.41 to 4.46 in surface soil and from 5.30 to 4.94 in subsurface soil across succession (Fig. 2A), whereas electrical conduc-tivity (EC) increased by 2.5-fold and 3.2-fold in surface and subsurface soils, respectively (Fig. 2F). Concentrations of NO_3_^−^-N and SO_4_^2−^ increased from 14.63 and 29.79 mg kg^−1^ in surface soil and from 19.02 and 15.82 mg kg^−1^ in subsurface soil at the 5-year stage to 120.05 and 56.42 mg kg^−1^ in surface soil and 90.70 and 33.88 mg kg^−1^ in subsurface soil at the 25-year and 40-year stages (Fig. 2H–I; Table S2). In contrast, five soil properties fluctuated: soil water content and carbon–nitrogen pools (TC, SOC and TN) peaked in mid-succession and then declined in both surface and subsurface soils, whereas the C:N ratio did not follow this shared pattern (Fig. 2B–E; Table S2). An exception was available phosphorus (AP), which showed no significant change among successional stages (Fig. 2G; Table S2). Together, these results indicate that edaphic recovery involved both directional trends and stage-specific fluctuations rather than a single linear trajectory.

### 3.2. Multilevel reassembly of soil bacterial and archaeal communities during succession

Different aspects of bacterial and archaeal communities showed distinct recovery trajectories. Bacterial and archaeal community composition varied across succession (db-RDA; bacteria: adjusted *R*^2^ = 0.357, *P* = 0.001; archaea: adjusted *R*^2^ = 0.330, *P* = 0.001), with fitted vectors linking the separation among stages to soil physicochemical gradients (Fig. 3A; Fig. S4A). These compositional differences among successional stages remained significant after accounting for elevation, soil depth, and the soil physicochemical variables included in the model (marginal PERMANOVA; bacteria: *R*^2^ = 0.064, *P* = 0.001; archaea: *R*^2^ = 0.058, *P* = 0.001; Fig. S3). The differences among stages were not accompanied by detectable differences in within-stage variability (betadisper; bacteria: *F* = 0.943, *P* = 0.421; archaea: *F* = 1.7736, *P* = 0.103). To assess whether compositional turnover represented recovery toward the primary-forest reference, we compared Bray–Curtis distances to the reference and found convergence in both bacterial and archaeal communities (Fig. 3B; Fig. S4B). Distances to the reference were positively associated with soil pH (linear mixed-effects models; bacteria: standardized *β* = 0.41, *P* < 0.001; archaea: standardized *β* = 0.33, *P* < 0.05; Fig. S5), indicating greater compositional similarity at lower soil pH.

**Figure 3:**
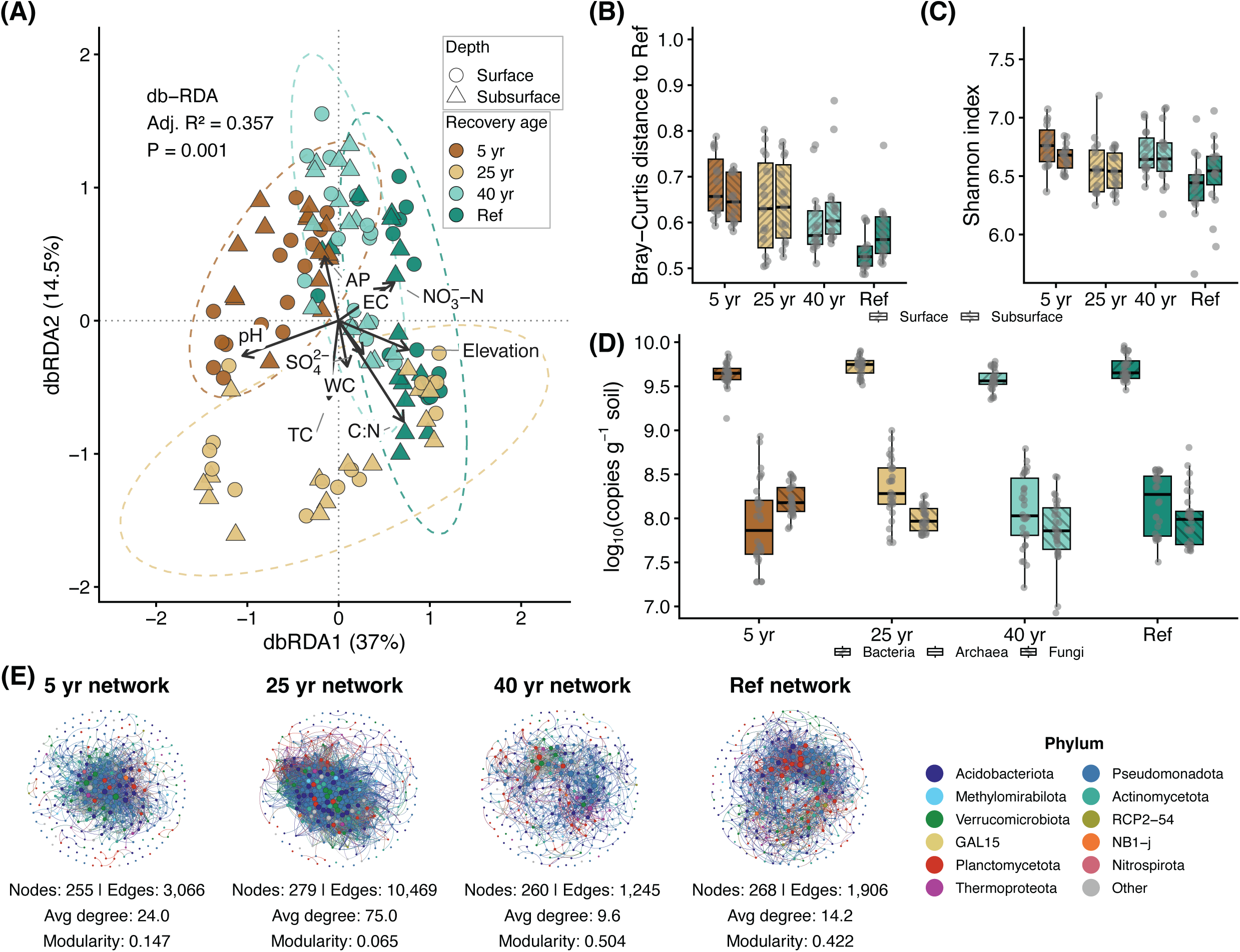
Multilevel reassembly of soil microbial communities across secondary forest succession. (A) Distance-based redundancy analysis (db-RDA) of bacterial community com-position. (B) Bray–Curtis distance to the primary-forest reference by soil depth. (C) Bacterial Shannon diversity by soil depth. (D) qPCR-based gene copy abundance, expressed as log_10_(copies g^−1^ dry soil). (E) Bacterial and archaeal co-occurrence networks; nodes are ASVs colored by phylum, and edges are significant associations between two ASVs. Node and edge counts, average degree, and modularity are shown for each network.

Compositional convergence was not accompanied by parallel recovery of Shannon diversity and qPCR-based abundance. Bacterial and archaeal Shannon diversity fluctuated across successional stages, while qPCR-based abundance showed group-specific patterns (Fig. 3C–D; Fig. S4C). Whereas compositional convergence was associated with soil pH, Shannon diversity in both bacteria and archaea was negatively associated with TC and SOC (linear mixed-effects models; bacteria: standardized *β* = -0.40 to -0.30; archaea: standard-ized *β* = -0.38 to -0.31; all *P* < 0.05; Fig. S5). Additionally, bacterial Shannon diversity was negatively associated with the soil C:N ratio (standardized *β* = -0.37, *P* < 0.01) and positively associated with pH (standardized *β* = 0.33, *P* < 0.05), and archaeal Shannon di-versity was negatively associated with TN (standardized *β* = -0.30, *P* < 0.05; Fig. S5). For qPCR-based abundance, bacterial and archaeal 16S rRNA gene abundances were positively associated with soil water content (bacteria: standardized *β* = 0.32, *P* < 0.01; archaea: stan-dardized *β* = 0.56, *P* < 0.001; Fig. S5). Fungal ITS abundance was negatively associated with AP (standardized *β* = -0.37, *P* < 0.01; Fig. S5).

Co-occurrence network organization also varied markedly across succession despite similar numbers of ASVs in each network (255–279; Fig. 3E). The 25-year network had the most co-occurrence associations per ASV (average degree = 75.0), compared with 24.0 at 5 years, 9.6 at 40 years, and 14.2 in the primary-forest reference. The 5 and 25-year networks also had low modularity (0.147 and 0.065, respectively), indicating less pronounced modular structure. In contrast, the 40-year and primary-forest networks combined lower connectivity with higher modularity (0.504 and 0.422, respectively), indicating fewer associations per ASV but more clearly defined modules (Fig. 3E). Thus, although connectivity and modularity varied non-monotonically across succession, the 40-year network resembled the primary-forest reference in its lower connectivity and higher modularity, contrasting with the more connected and less modular networks at 5 and 25 years. Together, these results show that community composition, Shannon diversity, qPCR-based abundance, and co-occurrence network organization did not recover in parallel.

### 3.3. Stage-specific reassembly of differentially abundant keystone taxa and their edaphic as-sociations

The dominant microbial phyla were consistent across succession, but their relative abundances varied. Bacterial communities were dominated by Acidobacteriota, Pseudomonadota, Verrucomicrobiota, GAL15, and Methylomirabilota (Fig. S6). Methylomirabilota was more abundant in the 5- and 25-year stages (11.8% and 9.48%, respectively), whereas Acidobacteriota and Pseudomonadota were enriched in the primary-forest reference (12.4% and 11.5%, respectively). Archaeal communities were dominated by Thermoproteota (53.6%–86.6%), followed by Thermoplasmatota (3.6%–15.0%); Halobacteriota was enriched in the 40-year stage (27.6%) (Fig. S6).

The similarity in network organization between the 40-year and primary-forest networks did not extend to the identity of putative keystone ASVs. Fifty-four ASVs were both differentially abundant across succession and classified as connectors or module hubs in stage-specific co-occurrence networks, with no ASV identified as a putative keystone in more than one stage (Fig. 4A). Their taxonomic composition also differed among stages: ASVs identified in the 5- and 25-year networks were mainly assigned to Acidobacteriota, Pseudomonadota, and Verrucomicrobiota, whereas Actinomycetota, Planctomycetota, and Methylomirabilota were additionally represented in the 40-year and primary-forest networks (Fig. 4A). ASVs identi-fied in the 5-year and primary-forest networks were generally less abundant in other stages, whereas those identified in the 25- and 40-year networks showed more variable cross-stage abundance patterns (Fig. 4B). These stage-specific sets also differed in their edaphic associations: early-stage ASVs were mainly associated with TN; ASVs identified in the 40-year network were negatively associated with SOC and TN but positively associated with NO^−^-N; and those identified in the primary-forest network showed fewer significant associations, mainly positive associations with SO^2−^ (Fig. 4C).

**Figure 4:**
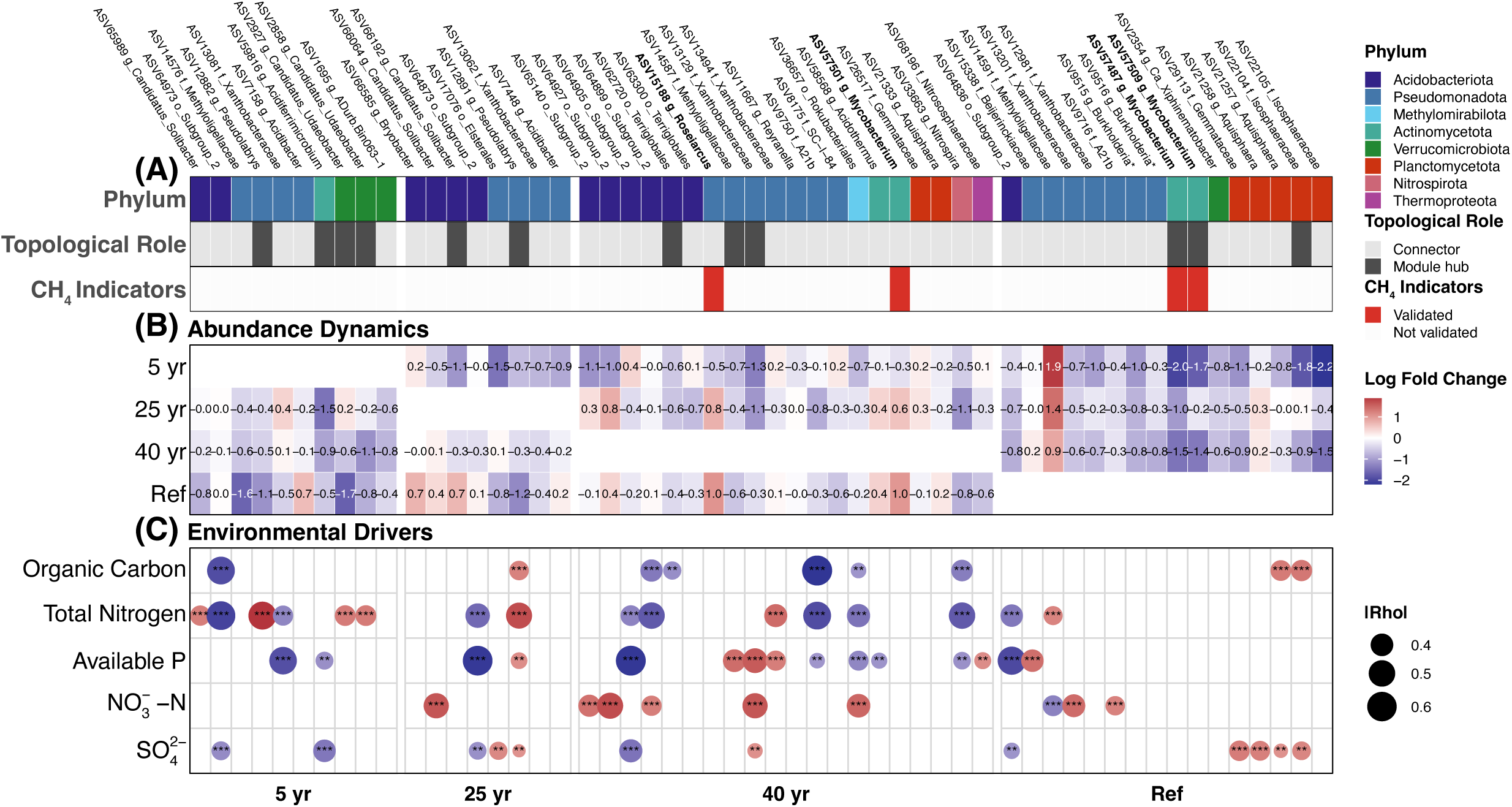
Abundance patterns and environmental drivers of differentially abundant key-stone ASVs across secondary forest succession in Metok. The 54 ASVs were identified by intersecting network topological roles with ANCOM-BC2 differential-abundance results and grouped by the stage in which each was identified as a keystone. (A) Phylum-level taxonomy, network topological role, and selection status in the exploratory Random Forest analysis of CH_4_ uptake. (B) ANCOM-BC2 bias-corrected log-fold changes, re-centered within each ASV relative to the stage in which it was identified as a putative keystone (red, higher; blue, lower). (C) Significant Spearman correlations between ASV abundance and soil variables; circles show associations with Benjamini–Hochberg-adjusted *P* < 0.05 and *|ρ| ≥* 0.30; size indicates *|ρ|* and color indicates direction. Asterisks denote adjusted *P* < 0.05 (*), < 0.01 (**), and < 0.001 (***).

### 3.4. Stage patterns of soil gas fluxes and microbial associations with methane uptake

In situ soil greenhouse-gas fluxes varied across succession (Fig. 5A). CH_4_ up-take was higher in the 25-year and primary-forest stands (mean fluxes: *−*2.683 and *−*2.773 nmol m^−2^ s^−1^) than in the 5- and 40-year stands (*−*1.094 and *−*0.899, respectively), with differences persisting after adjustment for elevation and WC (overall *P* = 0.0018; Holm-adjusted pairwise *P* = 0.0048–0.0137). In contrast, soil CO_2_ efflux showed weaker stage differentiation and a less stage-specific microbial signal (Fig. S7A–C). It was significantly higher in the 5- and 25-year stages than in the 40-year stage, whereas the primary-forest reference showed high within-stage variability and did not differ significantly from the secondary stages (Fig. S7A).

**Figure 5:**
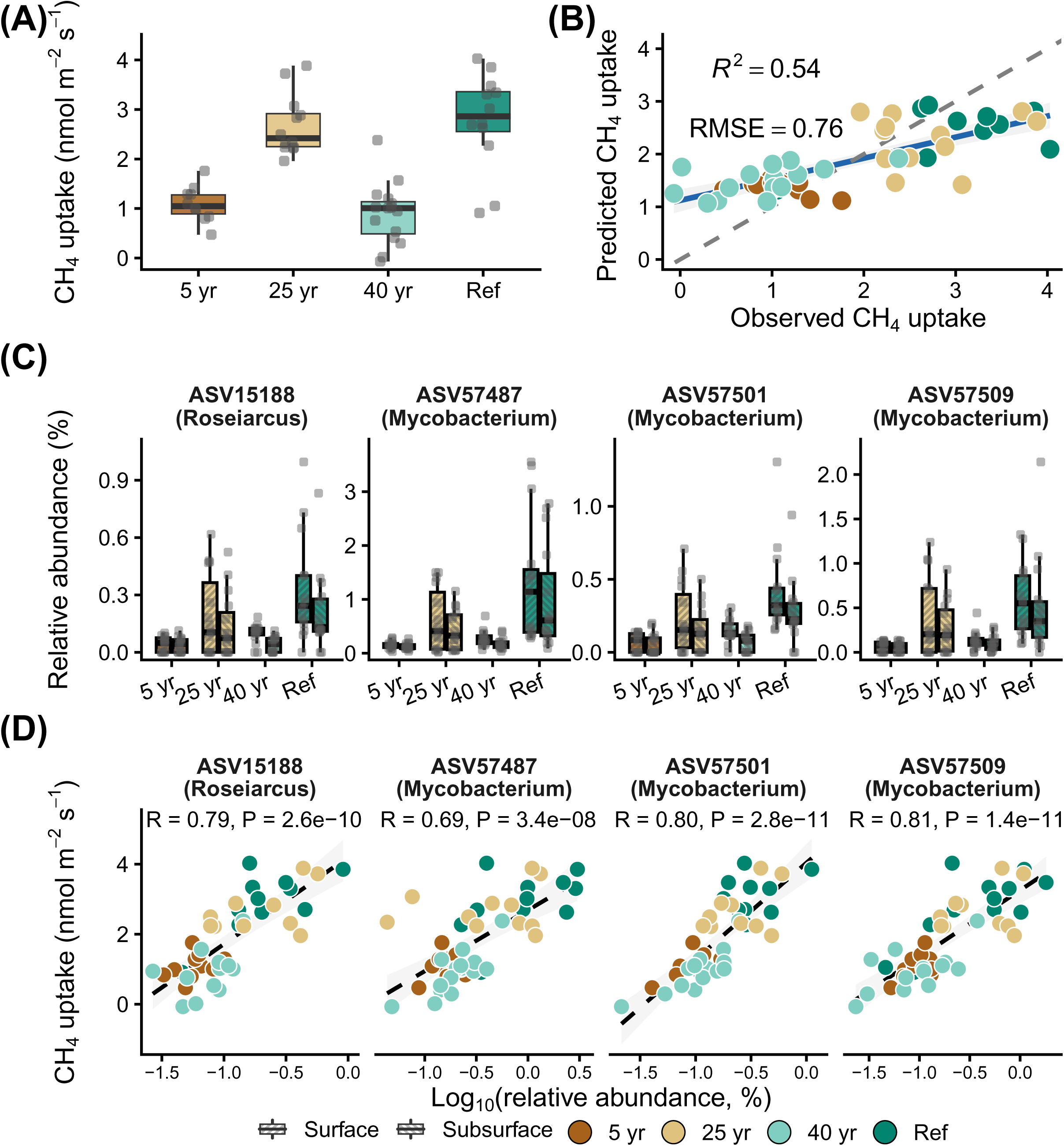
Soil CH_4_ uptake and exploratory associations with microbial ASVs across secondary forest succession. (A) CH_4_ uptake across successional stages. (B) Random Forest predictions of CH_4_ uptake from bacterial and archaeal community composition; the solid line shows the regression fit, shading indicates the 95% confidence interval, and the dashed line indicates the 1:1 relationship. (C) Relative abundances of the four selected ASVs by successional stage and soil depth. (D) Pearson correlations between log_10_-transformed ASV relative abundances and CH_4_ uptake; *R* and *P* values are shown. Boxes represent 25th–75th percentile of the data distribution with whiskers at 1.5 times the interquartile range and the middle line representing median.

Random Forest models identified the main predictors of greenhouse-gas fluxes. Soil physicochemical variables explained little variation in CH_4_ flux (*R*^2^ = 0.107; RMSE = 1.046; MAE = 0.842; Table S6), but more variation in CO_2_ flux (*R*^2^ = 0.197; RMSE = 0.952; MAE = 0.838; Table S7). TC, C:N ratio, soil organic carbon, and soil water content were the strongest predictors of CH_4_ flux (Table S6), whereas NO^−^-N was the strongest predictor of CO_2_ flux, followed by pH, electrical conductivity, and available phosphorus (Ta-ble S7). Bacterial and archaeal community composition explained more variation in CH_4_ flux (*R*^2^ = 0.54; RMSE = 0.76; MAE = 0.621; Fig. 5B) than in CO_2_ flux (*R*^2^ = 0.27; RMSE = 0.91; Fig. S7B). Putative methane-cycling taxa were characterized in the composite soil samples (Tables S3–S5). The taxonomically defined putative methanotroph set comprised 61 ASVs assigned to nine genera, with *Methylocapsa*, *Methylocystis*, and *Methylocella* accounting for 78.2%, 13.6%, and 6.9% of reads within this set across all soil samples, respectively. Their total relative abundances were 0.161%, 0.209%, 0.250%, and 0.314% in the 5-, 25-, and 40-year stages and the primary-forest reference, respectively (Tables S3–S4). Putative methanogenic archaea were much less abundant, with total relative abundances of 0.074%, 0.002%, 0.078%, and 0.032% in the 5-, 25-, and 40-year stages and the primary-forest reference, respectively. Their profiles were dominated by *Methanomassiliicoccales* (7.75%–62.54%) and *Methanomethyliaceae* (35.42%–90.34%) in the 5- and 25-year stages and the primary-forest reference, whereas *Methanosarcina* (54.09%) predominated in the 40-year stage.

The putative methanotroph set showed within-plot compositional associations with CH_4_ uptake, whereas its combined abundance provided limited additional predictive information. Combined relative abundance showed limited association with CH_4_ uptake after environmental adjustment and accounting for plot grouping (surface: *P* = 0.061; subsurface: *P* = 0.120). Estimated group-associated 16S copy abundance showed a positive surface-soil association (*P* = 0.030), but this did not meet the multiple-testing threshold (*q* = 0.059). Genus-level composition within the putative methanotroph set was described by the proportions of its nine genera relative to the set’s total read count. The first two principal components were jointly associated with CH_4_ uptake within plots after accounting for WC and the set’s combined relative abundance (surface: *P* = 0.038; subsurface: *P* = 0.027).

Complementary exploratory screening of the broader microbial community identified four ASVs through the intersection of random-forest ranking, differential abundance, and net-work position: three *Mycobacterium* ASVs and one *Roseiarcus* ASV (Fig. 5C). Their log_10_-transformed relative abundances showed positive correlations with CH_4_ uptake (*r* = 0.69–0.81; all *P <* 0.001; Fig. 5D). We compared the 16S rRNA gene sequences of the three *Mycobacterium* ASVs with those of previously reported methane-oxidizing *Mycobacterium* lineages to assess their similarity. The three *Mycobacterium* ASVs showed 97.87%–99.73% identity to published methane-oxidizing *Mycobacterium* references across 375–376 bp alignments (Van Spanning et al., 2022; Kambara et al., 2025). These associations remained conditional on selection from the same dataset, and the short 16S sequence similarities did not establish methane-oxidizing function.

## 4. Discussion

### 4.1. Directional recovery in soil inorganic properties and bacterial and archaeal community composition

Whether soil abiotic, biotic and functional recovery proceed synchronously during succession remains an important unresolved question. Our results from a subtropical forest chronosequence in the East Himalayas showed partial recovery of the soil ecosystem after traditional slash-and-burn disturbance. Specifically, soil pH, EC, nitrate, and sulfate, to-gether with bacterial and archaeal community composition, converged directionally toward the primary-forest reference (Fig. 2A,F,H–I; Fig. 3B). This pattern may reflect both the legacy effects of slash-and-burn agriculture on the soil environment and its subsequent reorganization during forest regeneration. The relatively alkaline but low-EC soil environment in the youngest forests was likely shaped by ash inputs together with slash-and-burn-induced losses of vegetation structure and canopy cover (Certini, 2005; Larssen et al., 2011; Veldkamp et al., 2020; Ke et al., 2021). As vegetation re-established, several coupled biogeochemical processes may have driven these soil properties toward the primary-forest reference. First, tree establishment likely promoted soil acidification by increasing plant uptake of base cations, including Ca^2+^, Mg^2+^, and K^+^. This process can deplete exchangeable base-cation pools in mineral soil and increase the relative dominance of H^+^ on exchange sites, as shown by a global afforestation meta-analysis (Berthrong et al., 2009) and a natural succession study (Jobbágy and Jackson, 2003). In addition, the humid climate and high precipitation in Metok may have intensified the leaching of exchangeable base cations, allowing older forests to experience greater cumulative base-cation losses and therefore lower soil pH (Yu et al., 2020). Second, vegetation recovery may have increased litterfall production and litter N concentration, thereby supplying more N-rich organic matter to the forest floor. This organic N can be mineralized to NH_4_^+^, increasing substrate availability for nitrification and promoting NO_3_^−^ formation (Yan et al., 2009). Enhanced nitrification may have further contributed to acidification, because the oxidation of NH_4_^+^ to NO_3_^−^ releases protons and can lower soil pH (Fujii et al., 2024). Third, recovery of canopy structure and root systems may have reduced hydrological losses while increasing organic S inputs to the soil, thereby helping to explain the increase in sulfate concentration during forest recovery (McGill and Cole, 1981; Scherer, 2009). Because EC reflects the total soluble ion strength of the soil extract, its increase is consistent with the parallel increases in NO_3_^−^ and SO_4_^2−^. Together, these processes offer plausible explanations for the observed differences in soil inorganic properties among successional stages.

In tropical and subtropical forests, these directional shifts in soil inorganic properties may have promoted bacterial and archaeal compositional convergence. Our results showed that convergence in bacterial and archaeal community composition was significantly associated with soil pH (Fig. S5). This relationship may arise because lower pH strengthens environmental filtering, which in turn promotes microbial community convergence. Strongly acidic conditions can lead to phylogenetically more clustered microbial communities through stronger deterministic assembly, whereas near-neutral pH conditions are associated with phylogenetically less clustered communities and greater stochasticity (Tripathi et al., 2018). As soils acidified during succession, bacterial and archaeal communities were likely assembled under stronger pH-mediated environmental filtering, selecting taxa adapted to acidic mature-forest conditions and thereby promoting bacterial and archaeal compositional convergence toward the primary-forest reference (Gubry-Rangin et al., 2011; Tripathi et al., 2013, 2018; Liu et al., 2021). This finding is consistent with previous studies across broad spatial scales (Lauber et al., 2009) and from tropical forest and cropland soil (Tripathi et al., 2013), where pH dominates variation in bacterial and archaeal community structure. In this sense, successional changes in pH likely promoted bacterial and archaeal compositional convergence by imposing acidity-related environmental filters that selected taxa adapted to mature-forest soil conditions.

### 4.2. Non-monotonic trajectories of soil organic resources, microbial diversity, abundance, and network structure

Based on reports that soil organic pools can recover to primary-forest levels within two decades (Martin et al., 2013; Veldkamp et al., 2020; Poorter et al., 2021), we expected soil carbon and nitrogen pools to recover rapidly and then stabilize. Instead, pools at 25 years were comparable to the primary-forest reference but declined markedly at 40 years (Fig. 2C–E), showing that intermediate similarity did not represent stable convergence. Long-term studies have attributed such non-monotonic patterns to mean annual temperature (Marín-Spiotta and Sharma, 2013), soil type (Paz et al., 2016), and forest functional composition (Wallwork et al., 2022, 2023). Because climate and soil type were broadly similar among our plots, shifts in forest functional composition provide a plausible explanation. Succession commonly replaces fast-growing pioneers with slower-growing mature-forest species (Wall-work et al., 2022). Pioneers supply abundant, nutrient-rich litter that decomposes rapidly (Poorter et al., 2004; Bakker et al., 2011; Szefer et al., 2017), whereas mature-forest species produce longer-lived, lower-quality litter (Kutsch et al., 2010). Pioneer establishment may therefore explain the larger organic pools and higher CO_2_ fluxes at 25 years, whereas declining high-quality litter inputs may account for their reduction at 40 years (Fig. 2C–E; Fig. S7) (Bakker et al., 2011; Wallwork et al., 2022). Greater root inputs in mature forests may further explain why organic pools were larger in the primary-forest reference than at 40 years (Veldkamp et al., 2020). Soil water content followed a similar stage pattern (Fig. 2B–E). Although it also depends on soil texture, bulk density, precipitation, and vegetation water use (Rawls et al., 2003; Seneviratne et al., 2010; Laio et al., 2001), soil organic carbon can increase porosity and water retention (Zhao et al., 2022). Thus, shifts in forest functional composition may have driven non-monotonic changes in soil organic matter and, indirectly, soil water content.

The non-monotonic patterns in Shannon diversity and qPCR-based abundance tracked different edaphic dimensions across succession (Fig. 3C–D; Fig. S4; Fig. S5). Both bacterial and archaeal Shannon diversity were negatively associated with TC and SOC. Bacterial diversity was additionally positively associated with pH and negatively associated with the C:N ratio, whereas archaeal diversity was additionally negatively associated with TN (Fig. S5). The positive relationship between bacterial diversity and pH is consistent with evidence that soil pH is a major predictor of bacterial alpha diversity (Fierer and Jackson, 2006; Lauber et al., 2009). High C:N conditions may intensify bacterial N limitation and stoichiometric filtering, reducing the number of taxa able to maintain growth (Zhang et al., 2016; Yang et al., 2022; Shang et al., 2021). For archaea, high N availability can favor dominant phyla and reduce diversity (Sun et al., 2019; Sadet-Bourgeteau et al., 2022); accordingly, high-TN samples showed greater Thermoproteota dominance and lower archaeal Shannon diversity (Fig. S5; Fig. S6). In contrast, bacterial and archaeal 16S rRNA gene abundances were positively associated with soil water content (Fig. S5), consistent with observations from Yunnan forests (Fu et al., 2020; Jia et al., 2024). Greater moisture may increase substrate diffusion and reduce osmotic stress, supporting larger microbial populations (Manzoni et al., 2012; Schimel, 2018). These processes may help explain the non-monotonic variation in diversity and abundance across stand ages.

Microbial co-occurrence network topology varied strongly among successional stages as soil organic resources and nutrient conditions shifted (Fig. 3E). Although higher soil organic carbon often supports greater network connectivity (Morriën et al., 2017; Wang et al., 2023), carbon quality can also shape network structure: recalcitrant organic matter may promote specialized, less connected, and more compartmentalized associations (Zheng et al., 2021; Hu et al., 2022a). The high connectivity of the 5- and 25-year networks, especially at 25 years, may therefore reflect greater labile-carbon availability, although carbon lability was not measured (Wallwork et al., 2023). In contrast, the lower connectivity and higher modularity at 40 years may reflect reduced or spatially heterogeneous substrate supply, whereas the similar topology of the primary-forest network may instead reflect a larger proportion of recalcitrant carbon (Wallwork et al., 2023; Xiong et al., 2021; Nie et al., 2024). Thus, topological similarity between the 40-year stage and the primary-forest reference may arise through different mechanisms. At 40 years, central taxa were negatively associated with organic carbon and total nitrogen but positively associated with nitrate (Fig. 4C), consistent with lower organic resources and stronger inorganic-N filtering, although these associations do not establish causality (Shao et al., 2019; Xin et al., 2022). In the primary-forest reference, central taxa were more stage specific and less strongly associated with bulk carbon and nitrogen pools (Fig. 4B–C), suggesting a greater role for unmeasured resource quality than bulk quantity (Bonner et al., 2020; Zeng et al., 2022). Network organization therefore remained stage specific rather than converging monotonically toward the primary-forest reference.

### 4.3. Environmental and microbial associations with soil CH_4_ uptake

Soil CH_4_ uptake varied among successional stages alongside differences in soil conditions and microbial composition. CH_4_ uptake was strongest in the 25-year stage and the primary-forest reference, but weaker in the 5-year and 40-year stages (Fig. 5A). While soil physicochemistry had a weak predictive power of soil CH_4_ sink function (*R*^2^=0.107), total carbon, C:N ratio, organic carbon, and soil water content were the best predictors of CH_4_ uptake (Fig. 2B–E; Table S6). Soil organic carbon can enhance forest CH_4_ sink strength by supporting methanotrophic activity (Lee et al., 2023). Soil moisture is also important: several studies have shown that CH_4_ often exhibits a U-shaped relationship with soil moisture (Castro et al., 1995; Dijkstra et al., 2011). Lower soil moisture may constrain microbial activity (Schimel, 2018), whereas high soil moisture can reduce O_2_ diffusion into the soil and create anaerobic microsites that favor methanogenesis (Le Mer and Roger, 2001; Conrad, 2020). In our study, higher SOC concentrations in the 25-year stage and the primary-forest reference may, on the one hand, have provided sufficient resources to support methanotrophic activity and, on the other hand, have improved soil structure and created a suitable moisture environment for methanotrophs.

The limited predictive contribution of combined abundance suggests that variation in CH_4_ uptake was not adequately captured by the quantity of taxonomically assigned putative methanotrophs alone. These DNA-based abundance measures neither quantify concurrent methane-oxidation activity nor provide complete coverage of methane oxidizers. In subtropical forests, drought enhanced CH_4_ uptake without changing methanotroph abundance (Zhou et al., 2021), while warming increased surface-soil uptake alongside reduced moisture and altered methanotroph composition, without a significant increase in *pmoA* copy abundance (Zhang et al., 2024). Such findings illustrate why microbial abundance and methane uptake need not respond in parallel to environmental change. In our study, the within-plot association between genus composition and CH_4_ uptake suggests that aggregation may obscure ecologically relevant differences among taxa. Similarly, nitrogen addition in a Moso bam-boo forest altered methanotroph composition and reduced CH_4_ uptake without significantly changing *pmoA* abundance (Li et al., 2021). These findings support considering composition alongside abundance when interpreting methane uptake.

Several *Mycobacterium*-affiliated ASVs were associated with CH_4_ uptake in the present dataset. The microbial Random Forest model yielded a higher *R*^2^ than the physicochemical model in the original analysis (Fig. 5; Table S6) and retained several *Mycobacterium*-affiliated ASVs among its top-ranked variables (Fig. S8). Three of the four selected ASVs belonged to this genus and were highly connected in the co-occurrence networks and positively correlated with CH_4_ uptake (Fig. 5C–D), although these network positions do not establish functional importance in methane cycling. Recent studies have demonstrated methane oxidation in particular *Mycobacterium* lineages (Van Spanning et al., 2022; Kambara et al., 2025), motivating further investigation of whether the taxa represented by our ASVs share this capacity. However, short 16S rRNA gene regions provide limited resolution among mycobacterial species (Tortoli, 2010; Walsh et al., 2019), and sequence similarity does not establish shared methane-oxidizing function. Alternative explanations should also be considered. Significant associations between *Mycobacterium*-affiliated taxa and methanotrophs have been reported in methane-amended peat (Kaupper et al., 2022). These observations provide a basis for considering indirect ecological links with methanotrophs. Alternatively, the association could arise from shared responses to soil conditions. Some *Mycobacterium*-affiliated ASVs showed lower relative abundances in compacted agricultural soils, alongside reduced air permeability and gas diffusion (Longepierre et al., 2021), suggesting that similar environmental sensitivities could link our taxa to conditions relevant to CH_4_ uptake. These explanations are not mutually exclusive and remain unresolved by the present observations. The CO_2_ results provide a complementary comparison. Soil CO_2_ efflux was only mod-estly predicted by bacterial and archaeal community composition in the original analysis, and no ASVs met all three screening criteria (Fig. S7). Both microbial composition and biomass have been associated with soil respiration rates and their responses to warming (Liu et al., 2018; Sáez-Sandino et al., 2023). Functional overlap documented across soil microbial communities provides a possible explanation for why taxonomic turnover did not translate into proportional changes in carbon-processing potential (Chen et al., 2022). These findings place the observed compositional associations within a broader context in which microbial identity, abundance and functional capacity contribute different information about soil processes. Considered alongside the CH_4_ stage patterns, the CO_2_ comparison highlights the complementary information provided by microbial community structure and soil gas fluxes.

## 5. Conclusions

This study reveals that belowground recovery after slash-and-burn disturbance in East-ern Himalayan secondary forests is a multidimensional and asynchronous process rather than a uniform return to the primary-forest reference. Directional changes in the soil inorganic environment were accompanied by bacterial and archaeal compositional convergence toward the primary-forest reference. However, other belowground properties, including soil organic pools, bacterial and archaeal Shannon diversity, 16S rRNA gene abundance, and co-occurrence network organization, showed fluctuating and stage-associated recovery patterns. Our inference is restricted to a space-for-time chronosequence in one montane subtropical forest region and to measurements made during a single seasonal observation window. Within this scope, atmospheric CH_4_ uptake showed a non-monotonic pattern across the chronosequence. The combined abundance of a taxonomically defined putative methanotroph set provided limited additional predictive information. Taxonomy composition within this set showed exploratory associations with CH_4_ uptake. Exploratory screening of the broader mi-crobial community also identified four ASVs—three assigned to *Mycobacterium* and one to *Roseiarcus*—whose relative abundances were positively associated with CH_4_ uptake. These findings extend previous assessments focused on individual soil properties and suggest that apparent abiotic, biotic or functional recovery at an intermediate stage may not be sustained over longer timescales. Longer chronosequences and repeated observations are therefore needed to evaluate both the extent and persistence of recovery across multiple belowground dimensions.

## CRediT authorship contribution statement

Mianhao Zhang: Conceptualization, Methodology, Software, Validation, Formal analysis, Investigation, Data curation, Writing – original draft, Visualization. Xiaohui Zhao: Methodology, Validation. Yuting Zhao: Methodology, Investigation. Dandan Shi: Investigation, Writing – original draft. Wenzheng Fu: Investigation, Writing – original draft. Cuiyun Yang: Investigation, Project administration. Hang Yu: Conceptualization, Methodology, Validation, Writing – review & editing, Supervision, Project administration.

## Data availability

The raw 16S rRNA gene amplicon sequencing reads have been deposited in the NCBI Sequence Read Archive (SRA) under BioProject accession number PRJNA1531034. The data on soil physicochemical properties, carbon fluxes, and prokaryotic communities supporting this study have been deposited in Zenodo (https://doi.org/10.5281/zenodo.23550284).

