## Supplementary information for "Decoupled recovery of soil physicochemistry, microbiome and methane sink function during secondary forest succession in the Eastern Himalayas"

### Supplementary Material

**Table S1.** Locations, elevations, and successional stages of the 12 sampling plots. Three plots represented each of the 5-year, 25-year, and 40-year secondary-forest stages and the primary-forest reference. Coordinates are given in decimal degrees and elevations in metres above sea level. Five sampling points were established in each plot, with soil collected at 0–10 and 10–20 cm depth at each point.

| Plot ID | Longitude | Latitude | Successional stage | Elevation |
|---|---|---|---|---|
| 25 yr-1 | 95.168˚E | 29.224˚N | 25-year | 1657 m |
| Ref-1 | 95.176˚E | 29.215˚N | Primary Forest | 1630 m |
| 5 yr-1 | 95.165˚E | 29.208˚N | 5-year | 1850 m |
| 5 yr-2 | 95.163˚E | 29.210˚N | 5-year | 1740 m |
| 40 yr-1 | 95.195˚E | 29.200˚N | 40-year | 1800 m |
| 40 yr-2 | 95.194˚E | 29.208˚N | 40-year | 1700 m |
| Ref-2 | 95.186˚E | 29.201˚N | Primary Forest | 1840 m |
| Ref-3 | 95.188˚E | 29.202˚N | Primary Forest | 1740 m |
| 25 yr-2 | 95.197˚E | 29.224˚N | 25-year | 1850 m |
| 40 yr-3 | 95.189˚E | 29.230˚N | 40-year | 1786 m |
| 25 yr-3 | 95.190˚E | 29.230˚N | 25-year | 1800 m |
| 5 yr-3 | 95.176˚E | 29.219˚N | 5-year | 1600 m |

**Table S2.** Soil physicochemical properties by successional stage and soil depth. Surface and subsurface soils correspond to 0–10 and 10–20 cm, respectively. Concentrations are expressed per unit dry soil. The C:N ratio is calculated for each sample as soil organic carbon divided by total nitrogen (SOC/TN) and is dimensionless. Values are means ± sample standard deviations (SD; n = 15 composite samples per stage and depth, from five sampling points in each of three independent plots). Within each property and depth, pairwise Wald z comparisons of estimated marginal means were obtained from the selected maximum-likelihood mixed-effects model used in Fig. S2, at mean elevation. P values were adjusted by the Holm method across the six stage comparisons within each property and depth. Groups sharing a lowercase letter are not significantly different (adjusted $P \geq 0.05$).

| Parameter | Depth | 5-year | 25-year | 40-year | Primary Forest |
|---|---|---|---|---|---|
| Total Carbon (g/kg) | Surface | 80.31 ± 20.36 b | 100.17 ± 21.34 a | 68.68 ± 9.50 b | 84.14 ± 23.31 b |
| | Subsurface | 58.16 ± 8.81 b | 70.61 ± 19.67 a | 48.71 ± 7.10 b | 57.67 ± 16.66 b |
| Total Nitrogen (g/kg) | Surface | 6.42 ± 1.89 a | 6.73 ± 1.40 a | 5.43 ± 0.82 a | 5.77 ± 1.99 a |
| | Subsurface | 4.31 ± 0.81 a | 4.83 ± 1.50 a | 3.60 ± 0.63 a | 3.83 ± 1.38 a |
| Organic Carbon (g/kg) | Surface | 74.35 ± 20.65 b | 94.51 ± 20.68 a | 63.35 ± 9.69 b | 79.37 ± 23.37 b |
| | Subsurface | 52.61 ± 9.24 b | 65.87 ± 17.90 a | 44.36 ± 7.82 b | 52.83 ± 16.40 b |
| C:N Ratio | Surface | 11.75 ± 1.40 a | 14.06 ± 0.99 a | 11.74 ± 1.33 a | 14.10 ± 2.16 a |
| | Subsurface | 12.28 ± 1.04 a | 13.81 ± 1.53 a | 12.39 ± 1.73 a | 14.23 ± 2.54 a |
| pH | Surface | 5.41 ± 0.55 a | 4.98 ± 0.93 ab | 4.56 ± 0.41 ab | 4.46 ± 0.64 b |
| | Subsurface | 5.30 ± 0.19 a | 4.96 ± 0.52 a | 5.00 ± 0.50 a | 4.94 ± 0.80 a |
| Water Content (%) | Surface | 36.18 ± 6.13 c | 41.66 ± 5.99 a | 37.97 ± 6.94 bc | 41.28 ± 5.89 ab |
| | Subsurface | 29.83 ± 6.33 c | 35.40 ± 5.16 a | 31.56 ± 3.73 bc | 33.87 ± 4.32 ab |
| Available P (mg/kg) | Surface | 7.70 ± 4.98 a | 7.45 ± 3.12 a | 12.53 ± 8.77 a | 8.06 ± 3.46 a |
| | Subsurface | 4.12 ± 1.94 a | 5.66 ± 3.99 a | 7.81 ± 5.99 a | 4.32 ± 2.28 a |
| Electrical Conductivity (µS/cm) | Surface | 104.05 ± 48.03 b | 201.24 ± 91.34 ab | 283.18 ± 136.88 a | 264.41 ± 150.64 a |
| | Subsurface | 54.73 ± 24.54 b | 113.30 ± 46.99 ab | 141.92 ± 77.59 ab | 174.59 ± 106.09 a |
| $NO_3^-$-N (mg/kg) | Surface | 14.63 ± 14.82 a | 54.60 ± 75.59 a | 120.05 ± 112.69 a | 98.19 ± 136.74 a |
| | Subsurface | 19.02 ± 17.82 a | 66.26 ± 88.73 a | 90.70 ± 78.15 a | 58.99 ± 65.00 a |
| $SO_4^{2-}$ (mg/kg) | Surface | 29.79 ± 14.43 b | 57.95 ± 43.10 a | 56.42 ± 27.04 ab | 59.14 ± 36.91 ab |
| | Subsurface | 15.82 ± 10.96 b | 39.88 ± 21.54 a | 33.88 ± 16.16 ab | 20.85 ± 10.73 ab |

**Table S3.** Total relative abundances of putative methane-oxidizing bacteria and methanogenic archaea across successional stages. Taxa were assigned to these groups using SILVA-based taxonomy, and abundances were calculated from the rarefied ASV table. Taxon-level contributions are listed in Tables S4 and S5, respectively. These assignments indicate potential methane-cycling capacity and do not confirm activity.

| Group | 5-year | 25-year | 40-year | Primary Forest |
|---|---|---|---|---|
| Putative methane-oxidizing bacteria | 1.787% | 1.188% | 1.135% | 0.880% |
| Putative methanogenic archaea | 0.074% | 0.002% | 0.078% | 0.032% |

**Table S4.** Relative abundances and within-group contributions of putative methane-oxidizing bacterial taxa across successional stages. Values outside parentheses give relative abundance in the analysed community; values in parentheses give each taxon's percentage of the total putative methane-oxidizing bacterial group within that stage. "Other taxa" combines the remaining assigned taxa. Taxonomy-based assignments do not establish methane oxidation by the detected ASVs.

| Taxon | 5-year | 25-year | 40-year | Primary forest |
|---|---|---|---|---|
| *Methylomirabilia* | 1.6117% (90.17%) | 0.9671% (81.41%) | 0.7865% (69.29%) | 0.5349% (60.77%) |
| *Methylocapsa* | 0.1321% (7.39%) | 0.1819% (15.31%) | 0.1109% (9.77%) | 0.2857% (32.46%) |
| *Methylocystis* | 0.0014% (0.08%) | 0.0095% (0.80%) | 0.1148% (10.11%) | 0.0012% (0.13%) |
| Others | 0.0422% (2.36%) | 0.0295% (2.48%) | 0.1229% (10.82%) | 0.0584% (6.64%) |

**Table S5.** Relative abundances and within-group contributions of putative methanogenic archaeal taxa across successional stages. Values outside parentheses give relative abundance in the analysed community; values in parentheses give each taxon's percentage of the total putative methanogenic archaeal group within that stage. "Other taxa" combines the remaining assigned taxa. Taxonomy-based assignments do not establish methane production by the detected ASVs.

| Taxon | 5-year | 25-year | 40-year | Primary forest |
|---|---|---|---|---|
| *Methanomassiliicoccales* | 0.0462% (62.54%) | 0.0007% (39.53%) | 0.0000% (0.00%) | 0.0025% (7.75%) |
| *Methanomethyliaceae* | 0.0262% (35.42%) | 0.0010% (55.81%) | 0.0166% (21.28%) | 0.0287% (90.34%) |
| *Methanosarcina* | 0.0003% (0.42%) | 0.0000% (0.00%) | 0.0422% (54.09%) | 0.0003% (0.89%) |
| Others | 0.0012% (1.62%) | 0.0001% (4.65%) | 0.0192% (24.63%) | 0.0003% (1.02%) |

**Table S6.** Predictive performance and variable importance of the random forest model for soil $CH_4$ flux using surface-soil physicochemical properties and elevation. Performance was evaluated by 50 repetitions of two-fold cross-validation grouped by plot. $R^2$ is the squared Pearson correlation between observed and pooled cross-validated predictions; RMSE and MAE are the root mean squared error and mean absolute error, respectively. Predictors are ranked by permutation importance from the full-data model. Selection frequency is the proportion of the 100 cross-validation fits in which a predictor ranked among the top ten. Mean importance is the average permutation importance over those fits in which that predictor ranked among the top ten. C:N denotes SOC/TN.

Model performance: $R^2 = 0.107$; RMSE = 1.046; MAE = 0.842

| Top predictors | Permutation importance | Selection frequency | Mean importance |
|---|---|---|---|
| C:N | 0.331 | 0.98 | 0.209 |
| Water content | 0.286 | 0.99 | 0.243 |
| Total carbon | 0.286 | 0.94 | 0.215 |
| Organic carbon | 0.286 | 0.96 | 0.211 |
| Total nitrogen | 0.0838 | 0.97 | 0.0901 |
| pH | 0.0827 | 0.89 | 0.0933 |
| Electrical conductivity | 0.0571 | 0.89 | 0.0506 |
| Available phosphorus | 0.0521 | 0.88 | 0.0750 |
| $SO_4^{2-}$ | 0.0513 | 0.91 | 0.0642 |
| $NO_3^-$-N | 0.0267 | 0.82 | 0.0930 |

**Table S7.** Predictive performance and variable importance of the random forest model for soil $CO_2$ flux using surface-soil physicochemical properties and elevation. Performance was evaluated by 50 repetitions of two-fold cross-validation grouped by plot. $R^2$ is the squared Pearson correlation between observed and pooled cross-validated predictions; RMSE and MAE are the root mean squared error and mean absolute error, respectively. Predictors are ranked by permutation importance from the full-data model. Selection frequency is the proportion of the 100 cross-validation fits in which a predictor ranked among the top ten. Mean importance is the average permutation importance over those fits in which that predictor ranked among the top ten. C:N denotes SOC/TN.

Model performance: $R^2 = 0.197$; RMSE = 0.952; MAE = 0.838

| Top predictors | Permutation importance | Selection frequency | Mean importance |
|---|---|---|---|
| $NO_3^-$-N | 0.667 | 1.00 | 0.257 |
| pH | 0.0859 | 0.97 | 0.159 |
| Electrical conductivity | 0.0400 | 0.94 | 0.0807 |
| Available phosphorus | 0.0369 | 1.00 | 0.0870 |
| Elevation | 0.0171 | 0.91 | 0.0728 |
| $SO_4^{2-}$ | 0.00685 | 0.87 | 0.0430 |
| C:N | 0.00674 | 0.92 | 0.0573 |
| Total carbon | 0.00350 | 0.86 | 0.0271 |
| Organic carbon | 0.00319 | 0.92 | 0.0264 |
| Water content | 0.00270 | 0.86 | 0.0221 |

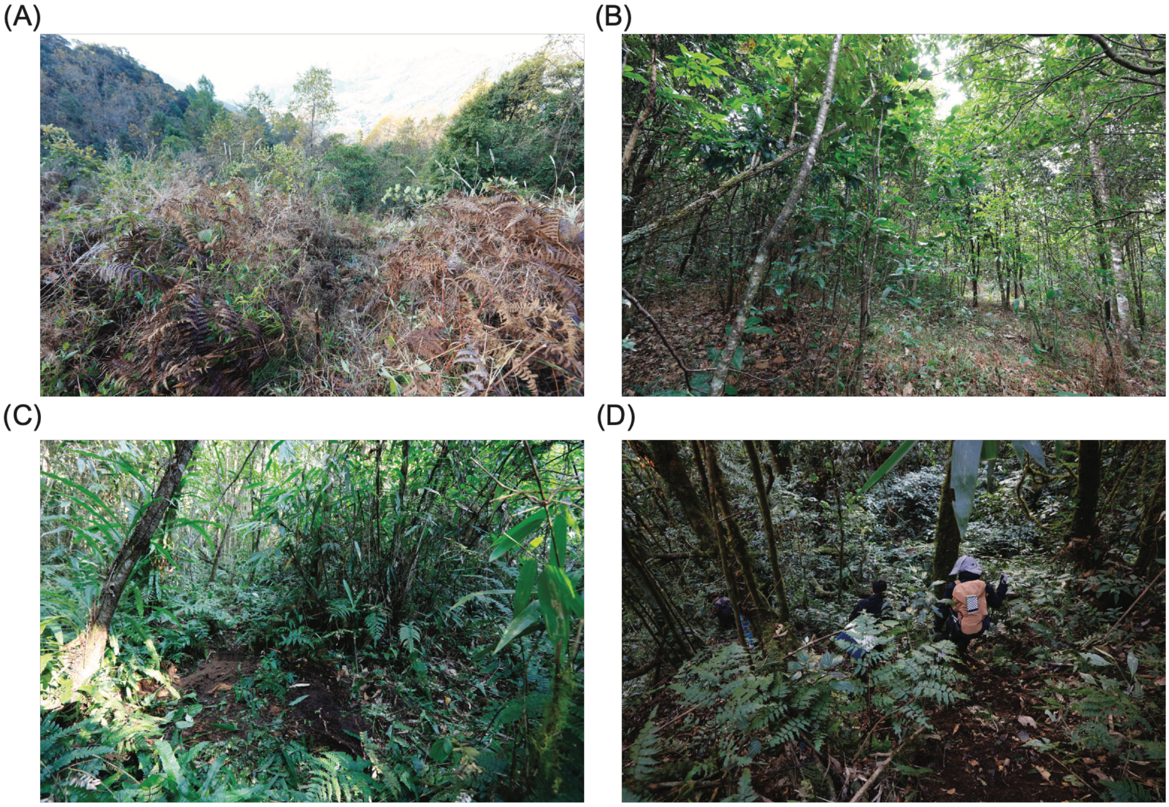


**Fig.** S1. Representative photographs of sampling plots across the forest chronosequence near Gelin Village, Metok County, Eastern Himalayas, China. (A) 5-year secondary forest; (B) 25-year secondary forest; (C) 40-year secondary forest; and (D) primary-forest reference.

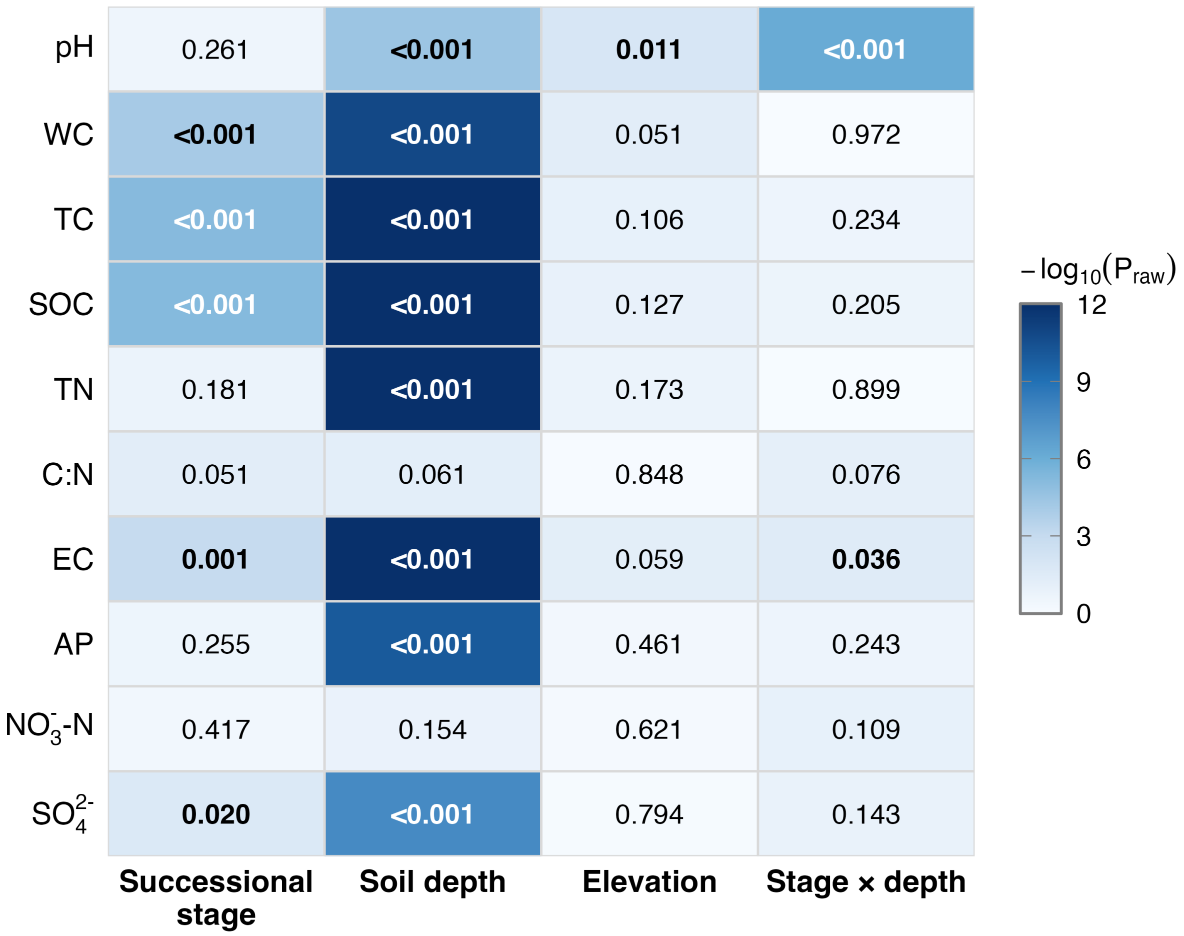


**Fig.** S2. Fixed-effect significance summary for soil physicochemical properties across secondary forest succession. Cell labels are raw omnibus P values from Type II Wald $\chi^2$ tests of linear mixed-effects models including successional stage, soil depth, elevation, and the successional stage × soil depth interaction as fixed effects, with random intercepts for sampling points nested within plots. The interaction was tested in the full model and removed when $P \geq 0.05$; main effects were then tested in the reduced additive model. Interaction P values always came from the full model. Bold values denote $P < 0.05$. Colour intensity represents $-\log_{10}(P)$, capped at 12, and indicates statistical evidence rather than effect size. Holm correction was applied only to post hoc pairwise stage comparisons. WC, water content; TC, total carbon; SOC, soil organic carbon; TN, total nitrogen; C:N, SOC/TN; EC, electrical conductivity; AP, available phosphorus.

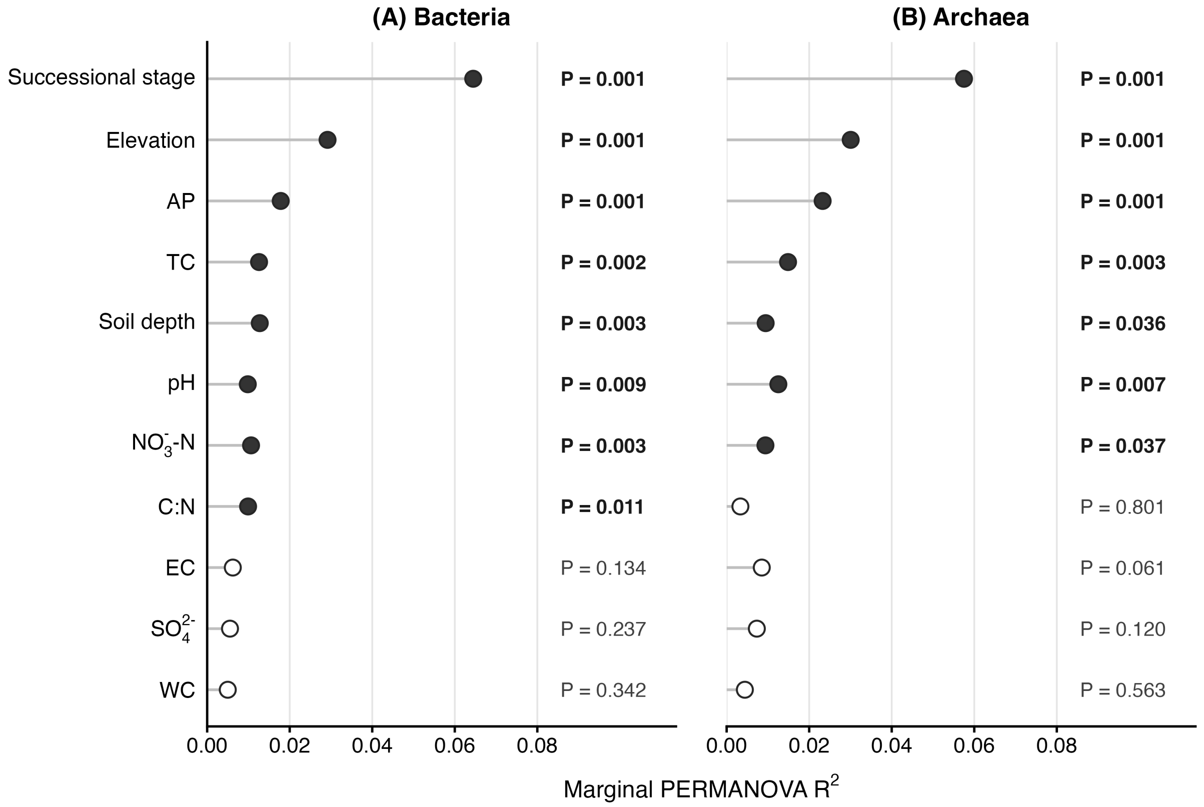


**Fig.** S3. Marginal PERMANOVA results for bacterial and archaeal community composition. Points show marginal $R^2$ values for successional stage, soil depth, elevation, and retained soil physicochemical variables. Horizontal lines extend from zero to the corresponding $R^2$. Filled and open points denote $P < 0.05$ and $P \geq 0.05$, respectively; labels give raw P values from 999 permutations. Soil predictors were screened for collinearity using a variance inflation factor threshold of 10. Spatial eigenvector covariates were included in the models but are omitted from the display. Retained predictors are shown in a common order based on the larger marginal $R^2$ across the two domains. WC, water content; TC, total carbon; C:N, SOC/TN; AP, available phosphorus; EC, electrical conductivity.

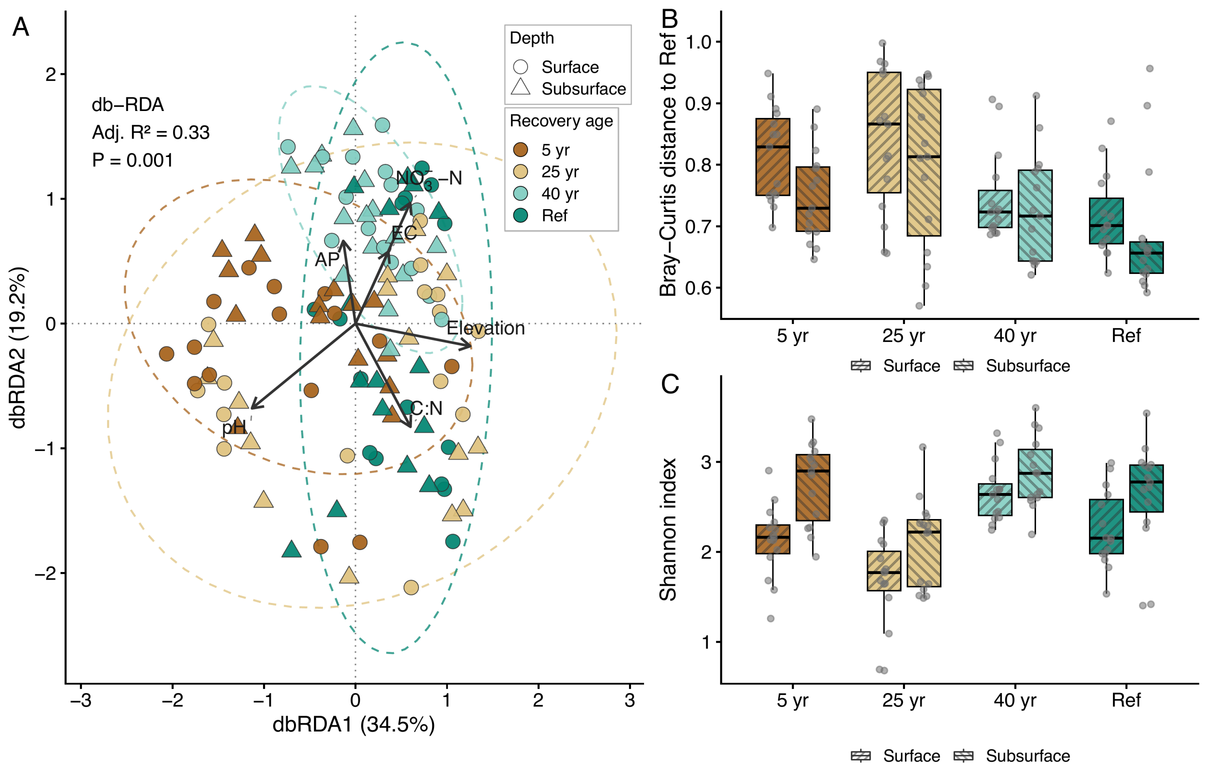


**Fig.** S4. Archaeal community composition and Shannon diversity across successional stages. (A) Distance-based redundancy analysis (db-RDA) of archaeal community composition; colours indicate stage and symbols indicate soil depth. Arrows show fitted environmental gradients. (B) Mean Bray–Curtis dissimilarity, calculated from ASV counts, to primary-forest samples at the same soil depth; reference samples were compared with the remaining reference samples at that depth. Lower values indicate greater similarity to the reference. (C) Archaeal Shannon diversity. Box centres and bounds indicate medians and interquartile ranges; points represent composite soil samples. Ref denotes primary forest.

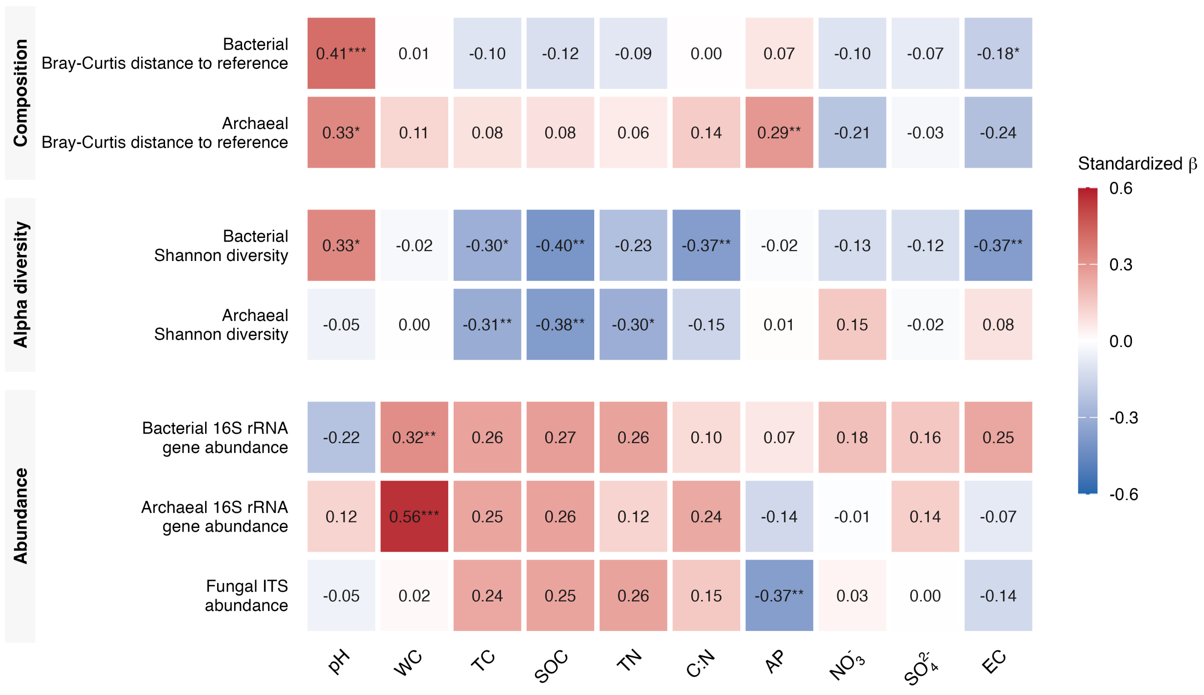


**Fig.** S5. Associations between microbial recovery metrics and soil physicochemical properties. Heatmap cells show standardized coefficients (β) from separate linear mixed-effects models for each microbial response–soil predictor pair. The seven responses comprise bacterial and archaeal Bray–Curtis distance to the primary-forest reference calculated from relative abundances, bacterial and archaeal Shannon diversity, $\log_{10}$ bacterial and archaeal 16S rRNA gene abundance, and $\log_{10}$ fungal ITS abundance. Microbial responses and soil predictors were standardized to z scores. Models included soil depth, successional stage, and elevation as fixed covariates, with random intercepts for sampling points nested within plots. Red and blue indicate positive and negative associations, respectively. Asterisks denote significance after Benjamini–Hochberg correction across all 70 response–predictor tests (*$P < 0.05$, **$P < 0.01$, ***$P < 0.001$). WC, water content; TC, total carbon; SOC, soil organic carbon; TN, total nitrogen; C:N, SOC/TN; AP, available phosphorus; EC, electrical conductivity.

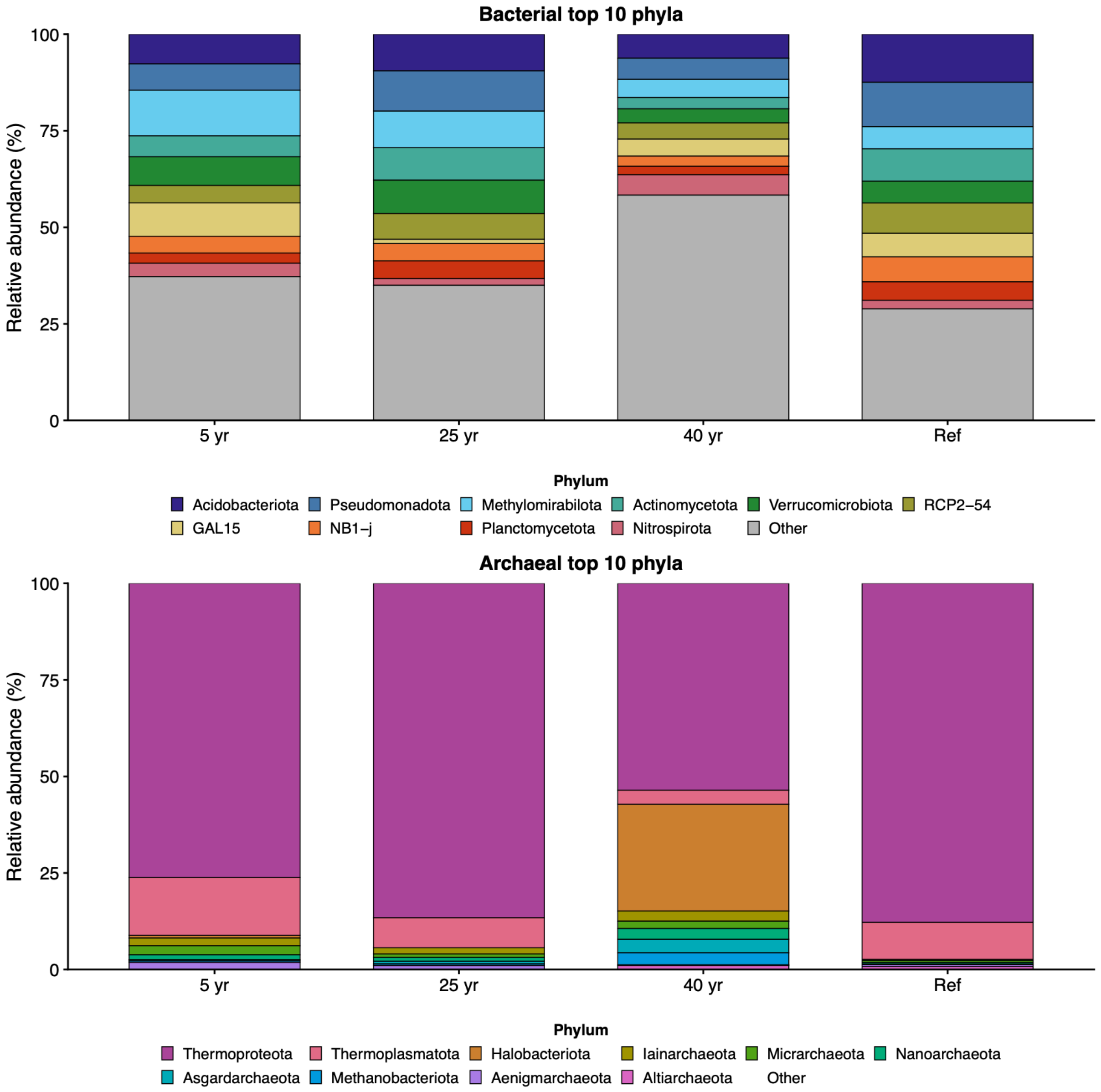


**Fig.** S6. Mean relative abundances of the ten most abundant bacterial phyla (upper panel) and archaeal phyla (lower panel) across successional stages. Bars show the 5-year, 25-year, and 40-year secondary forests and the primary-forest reference (Ref). "Other" includes the remaining phyla.

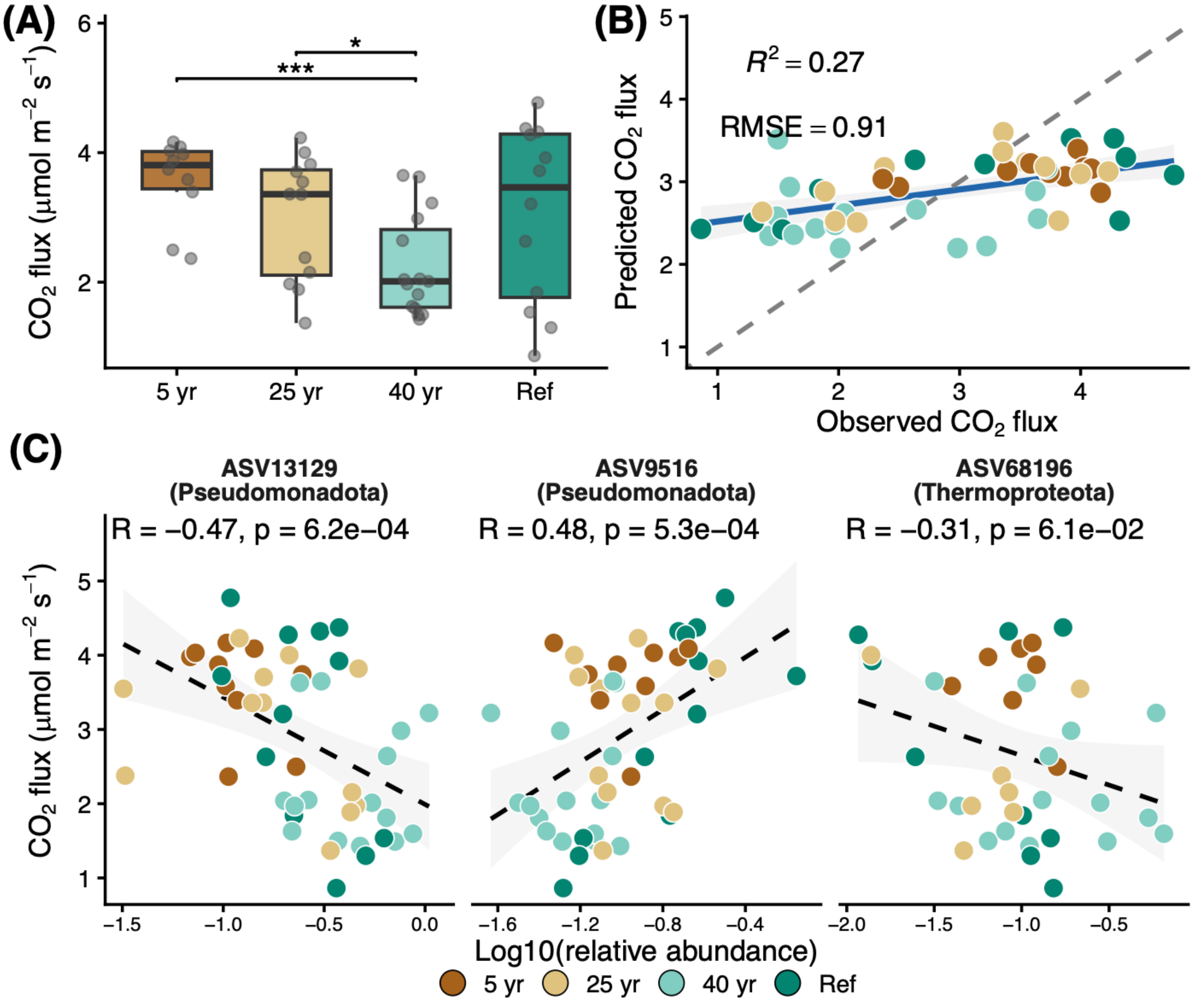


**Fig.** S7. Soil $CO_2$ flux across successional stages and its associations with microbial composition. (A) $CO_2$ flux in secondary forests and the primary-forest reference; positive values indicate soil $CO_2$ efflux. Box centres and bounds indicate medians and interquartile ranges. Brackets denote significant pairwise comparisons (* $P < 0.05$, *** $P < 0.001$). (B) Observed and random forest-predicted $CO_2$ flux based on microbial community composition; the grey dashed line marks the 1:1 relationship and the blue line shows the fitted trend. (C) Associations of $CO_2$ flux with $\log_{10}$-transformed relative abundances of ASV13129, ASV9516, and ASV68196, selected from the succession-associated keystone ASVs. Dashed lines show linear fits; Pearson R and P values are displayed. Colours identify successional stages, and Ref denotes primary forest.

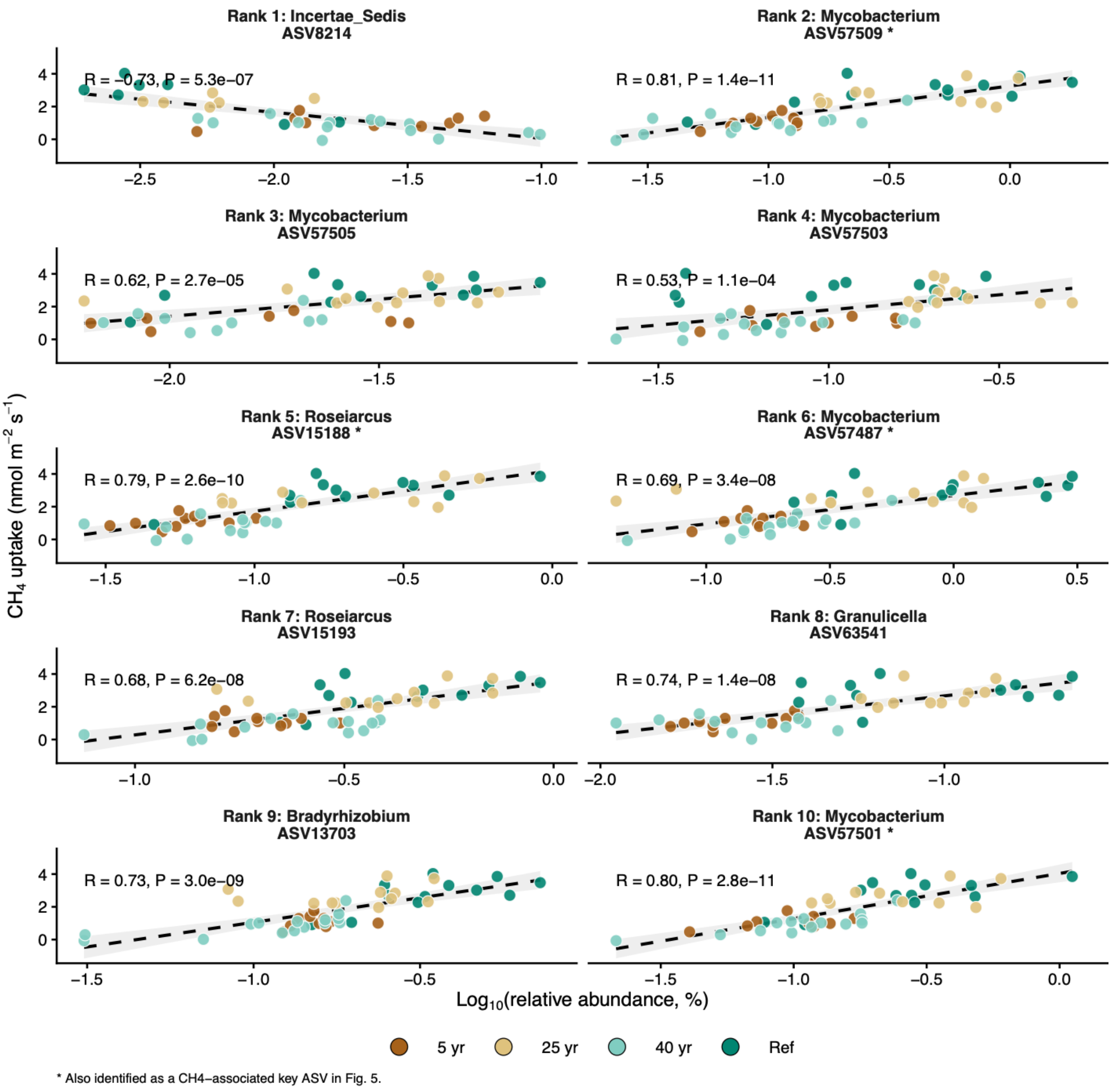


**Fig.** S8. Associations between the ten highest-ranked microbial random forest predictors and soil $CH_4$ uptake. Each panel relates $CH_4$ uptake to the $log_{10}$-transformed relative abundance of one ASV, expressed as a percentage before transformation. Points are coloured by successional stage; Ref denotes primary forest. Dashed lines show linear fits, and shading indicates 95% confidence intervals. Pearson R and P values are shown. Facet labels give predictor rank, ASV ID, and taxonomic annotation.
